# Beyond Abundance: Integrated Splicing and Expression Quantitative Trait Loci Refine Causal Gene Mapping in IBD

**DOI:** 10.64898/2026.11.22.768449

**Authors:** Emma Yu-Chen Huang, Shuying Han, Nolan H. Hamilton, Nina C. Nishiyama, Shehzad Sheikh, Terrence S. Furey

**Affiliations:** Department of Genetics, School of Medicine, The University of North Carolina at Chapel Hill, Chapel Hill, NC, USA; Curriculum in Bioinformatics and Computational Biology, The University of North Carolina at Chapel Hill, Chapel Hill, NC, USA; Center for Gastrointestinal Biology and Disease, School of Medicine, The University of North Carolina at Chapel Hill, Chapel Hill, NC, USA; Division of Gastroenterology and Hepatology, Department of Medicine, School of Medicine, The University of North Carolina at Chapel Hill, Chapel Hill, North Carolina, USA; Curriculum in Toxicology and Environmental Medicine, School of Medicine, The University of North Carolina at Chapel Hill, Chapel Hill, North Carolina, USA; Department of Biology, The University of North Carolina at Chapel Hill, Chapel Hill, NC, USA

**Keywords:** Splicing QTLs, Inflammatory Bowel Disease, Nonsense Mediated Decay, 5*^′^* untranslated region remodeling, protein domain alteration

## Abstract

Molecular quantitative trait loci for expression (eQTL) and splicing (sQTL) connect inflammatory bowel disease (IBD) risk loci to potential molecular mechanisms. We mapped *cis*-eQTL and *cis*-sQTL in 1,149 colon samples from SPARC IBD participants in IBD Plexus, identifying 20,566 genes with eQTL (eGenes) and 9,906 genes with sQTL (sGenes). Colocalization prioritized 239 eGenes and 164 sGenes across 147 distinct GWAS loci, including 58 genes supported by both traits. We assessed candidate splicing consequences for sQTL involving nonsense-mediated decay (NMD), 5*^′^* untranslated region modeling, and protein domain alteration. In five genes with GWAS-colocalized eQTL and sQTL, risk alleles were associated with reciprocal changes in total gene expression and the usage of junctions leading to predicted NMD-sensitive transcripts, consistent with NMD. IBD associated sGenes were enriched in immune and inflammatory pathways and included drug targets, some of which sQTLs may alter protein domains targeted by drugs, potentially contributing to variation in response to IBD therapies.

## Introduction

The inflammatory bowel diseases (IBD), primarily Crohn’s disease (CD), ulcerative colitis (UC), and indeterminate colitis (IC), are chronic inflammatory disorders of the gastrointestinal tract ^1^. Environmental triggers, gut microbiota, immune dysregulation, and host genetics all contribute to disease risk ^2,3^. Genome wide association studies (GWAS) have identified more than 300 IBD risk loci^4^. Most risk variants lie in non-coding regions, and their target genes and disease mechanisms often remain unclear^5^. Molecular quantitative trait loci (QTL) analyses help bridge this gap by linking genetic variants to molecular traits, such as gene expression or alternative splicing (AS), providing evidence for candidate target genes and generating hypotheses for mechanisms for how variants contribute to disease ^6^.

Expression QTLs (eQTLs) associate genetic variants with total gene expression levels. Recent disease bulk tissue colon eQTL studies collectively identified potential target genes for nearly 108 IBD loci^7^. While larger sample sizes and analyses in additional tissues may uncover eQTLs for some of the remaining loci, others may be driven by different mechanisms. One such mechanism is alternative splicing (AS), which allows one gene to produce multiple RNA isoforms, leading to, in some cases, protein isoforms with different functions ^8^. In IBD, alternative splicing has been linked to immune signaling, epithelial barrier function, cytokine regulation, and inflammatory responses^9,10^. Genetic variants that affect the frequency of using certain splice junctions, altering production of specific isoforms, are referred to as splicing QTLs (sQTLs).

For some genes, only an sQTL may be detected; thus, eQTL mapping may miss disease-related genetic mechanisms that act only through altered splice junction usage. For other genes both eQTL and sQTL may be present. These can result from variants primarily affecting gene expression, but in a way that disproportionately increases expression of some isoforms and decreases others, leading to the detection of an sQTL. However, splicing can also affect the detected abundance of an expressed gene. Certain splicing events can introduce a premature termination codon (PTC), leading to nonsense mediated decay (NMD) and reduced detected transcript abundance. Thus, sQTL that affect NMD frequency may also be detected as eQTL. In addition to NMD, sQTL can modulate the frequency of splicing events that affect the presence of certain translated protein domains or that can remodel the 5*^′^* untranslated region (UTR), which may affect transcript stability, translation efficiency, or regulatory binding^8^. Thus, combining sQTL with eQTL and GWAS colocalization analyses will more comprehensively elucidate how IBD-associated variants influence disease risk through both mRNA processing and gene expression.

To better characterize molecular effects of IBD risk variants, we mapped eQTL and sQTL in patient samples from the Crohn’s & Colitis Foundation IBD Plexus cohort. We colocalized these molecular QTLs with IBD GWAS loci signals to identify potential mechanisms contributing to disease risk. Our analyses supported many previously identified QTLs associated with IBD in colon tissue. In addition, we nominate potential target genes in 33 IBD GWAS loci lacking a QTL-based target and 174 disease prioritized genes across all loci that were not observed in other eQTL and sQTL colon tissue studies. We evaluated potential consequences of sQTL-associated splicing, including NMD, 5*^′^* UTR remodeling, and protein domain alteration. Five IBD risk genes, *ABO*, *ELMO1*, *GATD3*, *DR1*, and *YEATS4*, showed the combined colocalization and directional expression and splicing patterns expected for candidate NMD. In depth analysis of three identified risk genes, *PTPN2, TLR4,* and *ELMO1* show the diverse ways in which genetic variation may affect transcription and the functional roles of these genes in the colon. A review of therapeutic drugs for IBD and related diseases shows many target IBD eGenes and sGenes. We find some drugs target regions of proteins whose inclusion is affected by alternative splicing associated with sQTL. In total, these analyses connect IBD GWAS loci to candidate target genes that are involved in IBD-relevant biological pathways and provide potential RNA processing mechanisms that could contribute to increased risk.

## Results

### Large scale eQTL and sQTL discovery in the IBD Plexus patient cohort

We performed *cis*-eQTL and *cis*-sQTL analyses in 1,149 IBD colon tissue samples from the IBD Plexus cohort, in patient samples obtained from the Crohn’s & Colitis Foundation IBD Plexus program, specifically participants enrolled in the Study of a Prospective Adult Research Cohort with IBD (SPARC IBD); hereafter referred to as the IBD Plexus cohort (Fig. 1a). We identified eQTL for 20,566 genes (eGenes) with 18,246 distinct lead variants and sQTL for 43,131 spliced junctions (sJuncs) in 9,906 genes (sGenes) with 30,287 distinct junction-level lead variants and 9,297 distinct gene-level lead variants (FDR *<* 5%; Table 1, Fig. 1b; Supplementary Tables 1–4). We found that for 4,606 (46.5%) sGenes, there were multiple junctions with an sQTL, with an average of 4.35 significant junctions (Supplementary Fig. 1). As junction level lead sQTLs retained the identity of the associated splice event and supported more precise mechanistic interpretation than gene level lead sQTLs, we focused on junction level sQTLs in downstream analyses (Supplementary Figs. 2–3; Supplementary Results). These eQTL and sQTL strongly replicated colon QTL found by GTEx^11^, BarcUVa ^12^, and our previous independent IBD cohort study (UNC)^7^ (replication rates eQTL 0.879-0.99; sQTL 0.948-0.96; Fig. 1c; Table 2; Supplementary Tables 5–6). Cross cohort effect estimates also showed high concordance, particularly for sQTLs aligned to the same junction (Supplementary Figs. 4–5).

**Figure 1:**
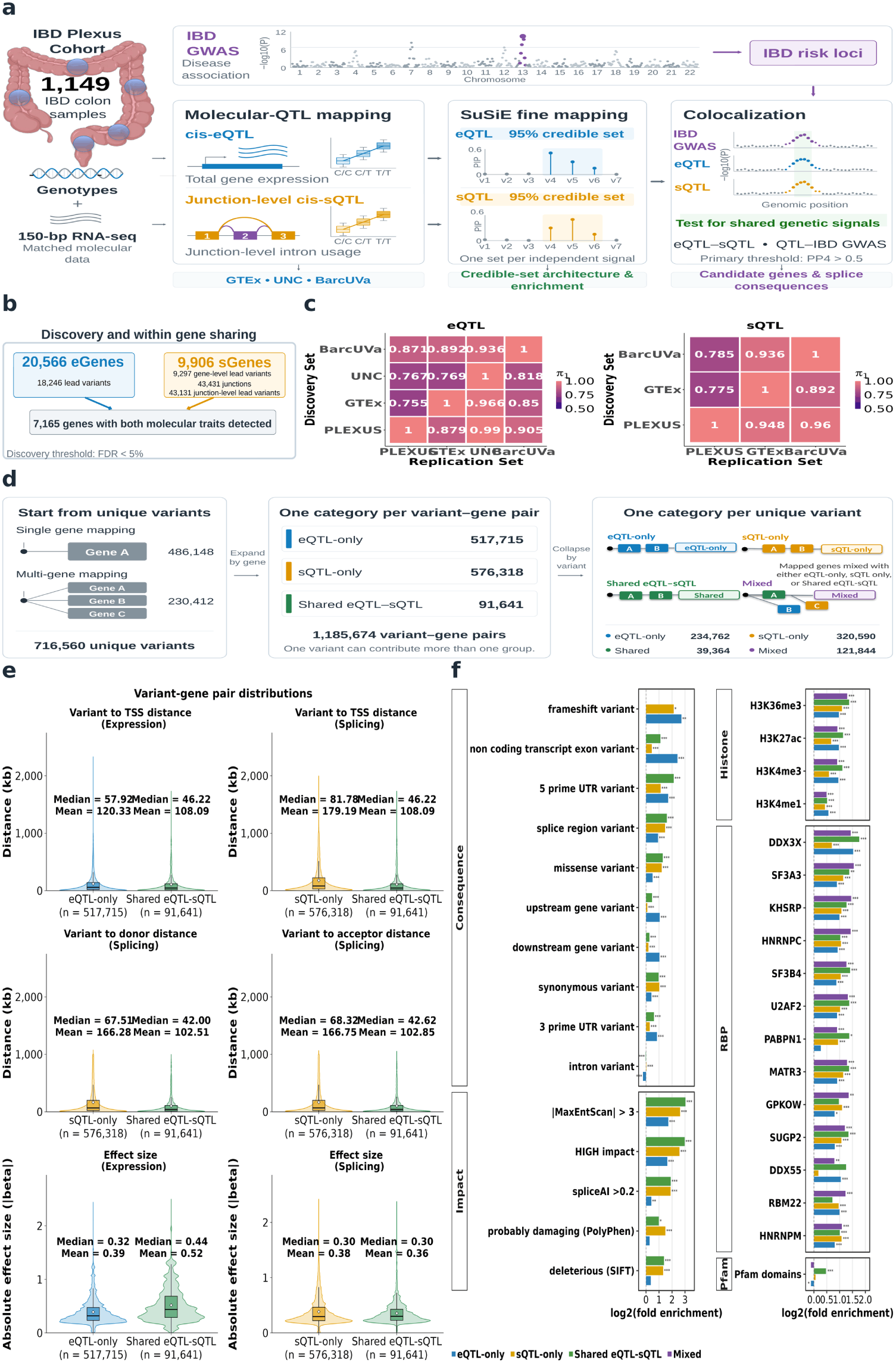
Study design and the IBD Plexus molecular QTL characteristics. **(a)** Workflow to perform eQTL and sQTL analyses and colocalization with IBD GWAS using colon biopsies from 1,149 IBD Plexus patients. **(b)**Detected eQTL and sQTL. **(c)**Directional cross-cohort *π*_1_ replication of eQTL and sQTL associations among IBD Plexus, GTEx, UNC, and BarcUVa colon cohorts. **(d)** Workflow and categorization of results from variant credible-set analysis. **(e)** Distributions of distances between variant-to-transcription-start-site (TSS), variant-to-splice-donor, and variant-to-splice-acceptor, and of absolute eQTL and sQTL effect sizes for trait-specific and shared variant–gene pairs. Embedded box plots show medians and means. **(f)** Functional enrichment of credible-set variant–gene pairs across variant consequences and predicted impact categories (left), and of unique credible-set variants across histone marks and RNA-binding protein (RBP) sites (right). Horizontal bars show log_2_(fold enrichment) relative to the matched background, colored by QTL-sharing category. Asterisks indicate statistical significance: *^∗^*:adjusted *P*: *<* 0.05;*^∗∗^*:adjusted *P <* 0.01;*^∗∗∗^*: adjusted *P <* 0.001.

**Table 1:** Summary of sQTL and eQTL discoveries in the IBD Plexus cohort.

| **Type of QTL** | **Number of Tested Genes (Junctions)** | **FDR 5% (p-value)** | **Number of Significant Genes (Junctions)** | **Unique Lead SNPs per Gene (Junctions)** | ***cis*-QTLs** |
|---|---|---|---|---|---|
| sQTL | 15,030 (201,258) | 5.64 × 10<sup>−3</sup> | 9,906 (43,131) | 9,297 (30,287) | 1,337,917 |
| eQTL | 41,005 | 2.39 × 10<sup>−2</sup> | 20,566 | 18,246 | 1,569,755 |

*Notes.* Summary of cis-sQTL and cis-eQTL discoveries in the IBD Plexus cohort. Numbers in parentheses indicate junction level results for sQTL analyses. The reported FDR thresholds correspond to the nominal p-value cutoffs controlling the false discovery rate at 5%.

**Table 2:** Summary of eQTL and sQTL discoveries across cohort.

| **Dataset** | **Sample Size** | **Tested Variants** | **eGenes** | **Lead eQTLs** | **sGenes** | **Gene-level**<br>**Lead sQTLs** |
|---|---|---|---|---|---|---|
| IBD Plexus | 1,149 | 9.1M (MAF ≥ 0.01) | 20,566 | 18,246 | 9,906 | 9,297 |
| GTEx v8 | 368 | 10.0M (MAF ≥ 0.01) | 11,686 | 11,303 | 3,459 | 3,306 |
| BarcUVa | 445 | 6.8M (MAF ≥ 0.01) | 11,737 | 11,425 | 1,062 | 1,059 |
| UNC | 252 | 8.4M (MAF ≥ 0.02) | 4,087 | 3,802 | – | – |

*Notes.* Summary of eQTL and sQTL discoveries in IBD Plexus, GTEx v8, BarcUVa, and UNC at a false discovery rate of 5%. Significant genes represent unique eGenes or sGenes associated with at least one significant variant. Lead eQTLs and gene level lead sQTLs represent unique lead variants at the gene level. MAF, minor allele frequency.

We performed fine mapping to identify credible sets, resulting in 713,842 unique sQTL and 609,356 unique eQTL variants corresponding to 1,185,674 unique variant–gene pairs. To better understand the characteristics of these variants, we first partitioned unique variant-gene pairs into eQTL-only (517,715 variants), sQTL-only (576,318 variants), and shared eQTL–sQTL (91,641 variants) groups (Fig. 1d). Because a variant may map to a single or multiple genes, each variant–gene pair was classified independently into one of these three groups according to its association with the paired gene. Shared eQTL–sQTL variants were closer to transcriptional start sites (TSSs; median 46.2 kb) than eQTL-only (median 57.9 kb) and sQTL-only (median 81.8 kb) variants. Shared eQTL–sQTL variants were also closer to splicing donor and acceptor sites than sQTL-only variants (donor: median 42.0 kb versus 67.5 kb; acceptor: 42.6 kb versus 68.3 kb) (Fig. 1e). Interestingly, the shared eQTL-sQTL had significantly stronger expression-related effect sizes than for eQTL-only, while sQTL effect sizes were similar across categories. These indicate that shared eQTL–sQTL variants preferentially localize near genomic features where transcriptional and splicing regulation may converge and tend to affect expression more strongly. Effects on both expression and splicing suggest expression of specific isoforms for a gene may vary, with some transcripts increasing or decreasing proportionally more than others.

We further compared genomic annotations and predicted functional consequences across the three variantgene pair categories, including high impact consequences annotated by the Ensembl Variant Effect Predictor (VEP)^13^, such as frameshifts, gained stop codons, or alterations to splice donor or acceptor sites. We assessed enrichment using variant–gene pairs classified as neither eQTLs nor sQTLs and matched for allele frequency and genomic region as the background set (Fig. 1f; Supplementary Tables 7–8). We found that eQTL-only variants were enriched in exons within non-coding transcripts (5.3-fold, adjusted *P* = 6.12 *×* 10^−308^) and frameshift variants (6.6-fold, adjusted *P* = 0.000738), while sQTL-only variants were enriched in spliced regions (2.76-fold, adjusted *P* = 4.99 *×* 10^−41^) and intronic regions (1.02-fold, adjusted *P* = 5.50 *×* 10^−24^). sQTL-only variants also showed higher proportions of predicted splice altering variants (3.70-fold, adjusted *P* = 2.55 *×* 10^−97^; SpliceAI ^14,15^ scores *>* 0.2), missense variants classified as deleterious by SIFT^16^ (2.50-fold, adjusted *P* = 9.11 *×* 10^−24^), and missense variants classified as probably damaging by PolyPhen (2.81-fold, adjusted *P* = 7.41 *×* 10^−16^)^17^. Shared eQTL–sQTL variants showed both patterns, with prominent enrichment in 5*^′^* untranslated regions (UTRs) (4.37-fold, adjusted *P* = 1.67 *×* 10^−31^) and spliced regions (3.04-fold, adjusted *P* = 2.46 *×* 10^−12^), as well as enrichment of predicted splice altering variants by SpliceAI (3.73-fold, adjusted *P* = 1.91 *×* 10^−22^) and SIFT-deleterious missense variants (2.6-fold, adjusted *P* = 1.56 *×* 10*^−^*^6^). All three groups were enriched for variants with large predicted changes in splice site strength, as assessed by MaxEntScan (eQTL-only: 3.27-fold, adjusted *P* = 1.14 *×* 10^−14^; sQTL-only: 6.16-fold, adjusted *P* = 3.51 *×* 10^−70^; Shared eQTL–sQTL: 8.19-fold, adjusted *P* = 7.8 *×* 10^−25^)^18^.

To explore potential regulatory mechanisms underlying the different QTL groups, we examined their overlap with histone marks from the ENCODE project, RNA-binding protein (RBP) binding sites from SKIPPER, and genomic regions encoding protein domains from the UCSC Genome Browser (Supplementary Methods). We compared four categories: eQTL-only, sQTL-only, shared eQTL–sQTL, and mixed-QTL variants (Fig. 1f; Supplementary Tables 9–10). The mixed-QTL group comprised multi-gene variants for which both eQTL and sQTL were detected but associated with different genes, whereas the eQTL-sQTL variants were associated with the same gene (Fig. 1d). As expected, eQTL-only variants showed greater enrichment for histone marks corresponding to active promoters and enhancers (H3K4me3, H3K4me1, and H3K27ac; 1.48–1.96-fold versus 1.35–1.59-fold; all adjusted *P <* 4.42 *×* 10^−144^), while sQTL-only variants showed greater enrichment for marks associated with splicing (H3K36me3; 1.96-fold versus 2.13-fold; adjusted *P <* 1.34 *×* 10^−307^). Shared eQTL–sQTL variants showed a combination of these features with strong enrichment for active gene regulatory histone marks (H3K4me3, H3K4me1, H3K27ac, H3K36me3; 1.42–2.58-fold; all *P <* 9.27 *×* 10^−15^) as well as RBP binding sites for spliceosomal factors such as SF3B4 (2.64-fold; *P <* 2.28 *×* 10^−12^). These eQTL-sQTL variants also showed the greatest overlap with gene regions encoding functional protein domains identified by Pfam (1.38-fold; *P <* 1.81 *×* 10*^−^*^4^), further supporting potential effects on protein function. Similar to shared eQTL-sQTL, mixed-QTL were enriched in most of the active regulatory histone marks and RBP binding sites, which of note they showed the greatest overlap with RBPs (HNRNPC: 2.69-fold, *P* = 1.07 *×* 10^−52^ and GPKOW: 2.55-fold, *P* = 1.39 *×* 10*^−^*^3^), but were not enriched in gene regions encoding functional protein domains.

### IBD GWAS colocalization shows locus complexity and identifies new potential gene targets

To identify potential IBD risk genes, we colocalized our eQTL and junction level sQTL signals with 320 IBD GWAS loci^4^. We found 239 colocalizing eQTLs (GWAS eGenes) across 121 unique IBD loci and 610 colocalizing sQTL signals in 164 sGenes (GWAS sGenes) across 97 unique IBD loci (posterior probability of same genetic signal (PP4)*>* 0.5; Supplementary Tables 11–12). The union comprised 345 genes (GWAS colocalized) across 147 GWAS loci. Loci contained a mixture of eQTL and/or sQTL, with 50 having only GWAS-colocalized eQTL (eQTL-only loci), 26 having only GWAS-colocalized sQTL (sQTL-only loci), and 71 having both (shared eQTL-sQTL loci) (Fig. 2a).

**Figure 2:**
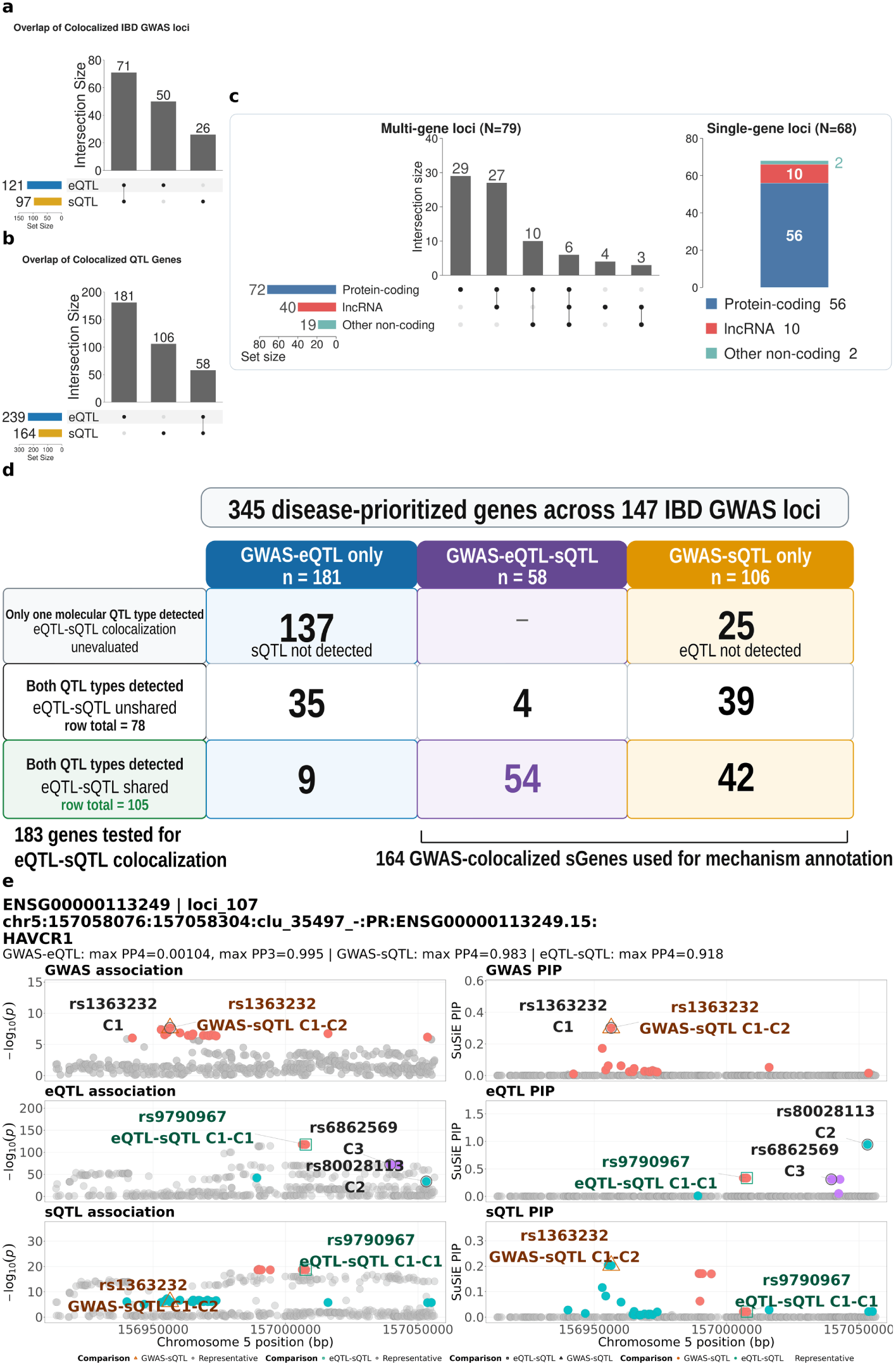
IBD GWAS colocalization and disease-prioritized gene classes. **(a)** Of 147 IBD GWAS loci, numbers that contain colocalized eQTL only, sQTL only, or both. The two QTL types at a shared locus could implicate different genes. **(b)** Detected QTL types for 345 genes colocalized with IBD GWAS. **(c)** Transcript types of genes in 79 multi-gene and 68 single-gene GWAS loci. **(d)** Classification of 345 disease-prioritized genes across GWAS-sharing status, type of QTL detected, and within-gene eQTL–sQTL colocalization. **(e)** Pairwise colocalizations for *HAVCR1* gene. Regional association statistics (*−* log_10_ *P*, left) and SuSiE posterior inclusion probabilities (PIPs, right) are shown for IBD GWAS, gene expression, and splice junction usage (top to bottom). Two components of the same sQTL colocalized with different traits. sQTL component 2 colocalized with GWAS component 1 (PP4 = 0.983), with rs1363232 highlighted as a shared candidate variant (triangle). sQTL component 1 colocalized with eQTL component 1 (PP4 = 0.918), with rs9790967 highlighted as a shared candidate variant (square). The direct GWAS–eQTL comparison favored distinct genetic associations (PP3 = 0.995).

We categorized GWAS colocalized genes based on whether they had a colocalized eQTL, a colocalized sQTL, or both (GWAS colocalization sharing status). For 58 genes, we detected both eQTL (24.2% of GWAS eGenes) and sQTL (35.4% of GWAS sGenes) that colocalized to the GWAS signal (GWAS eQTL-sQTL), while for 181 genes only eQTL (75.8%) colocalized (GWAS eQTL-only) and for 106 genes only sQTL (64.6%) colocalized (GWAS sQTL-only) (Fig. 2d; Supplementary Table 13). Note that GWAS eQTL-only and GWAS sQTL-only genes can fall within shared eQTL-sQTL loci if they target distinct genes. Interestingly, there were 125 genes for which eQTL and sQTL were detected, but for which only one of the QTL colocalized with a GWAS signal. In total, 108 GWAS eQTL-only, 77 GWAS sQTL-only, and 58 GWAS eQTL-sQTL genes fell within the 71 shared eQTL-sQTL loci.

Of the 147 colocalized IBD risk loci, 79 (53.7%) were associated with QTL for more than one gene. To understand what types of genes contributed to this complexity, we examined the gene biotypes at these loci. Most multi-gene loci included a QTL for at least one protein coding gene (72 of 79, 91.1%), and the most common pattern was QTL for multiple protein coding genes without a noncoding gene QTL (29 loci). We found that 40 loci included at least one QTL for a long non-coding RNA (lncRNA), of which 33 also included a QTL to a protein-coding gene. Nineteen included at least one QTL for a gene assigned to another noncoding gene biotype, all of which also had a QTL for a protein coding gene, a lncRNA, or both (Fig. 2c). Non-coding RNAs have been shown to have diverse roles in gene regulation, especially for proximal genes^19^. Thus, while these patterns may simply reflect shared genetic regulation of nearby coding and non-coding genes, there may also be regulatory relationships between the non-coding RNAs and the coding genes mediating the QTL signal. Together, these indicate that for most IBD GWAS loci, there may be multiple genes that contribute to genetic risk, some indirectly, highlighting the need to consider all genes supported by colocalization rather than assuming the presence of a single candidate.

Compared with previous colon tissue QTL studies by GTEx^11^, BarcUVa^12^, and UNC^7^, our analysis uniquely identified targets for 33 GWAS loci and QTL for 174 colocalized genes (108 GWAS eQTL-only genes, 58 GWAS sQTL-only genes, and 8 GWAS eQTL-sQTL genes) in 102 GWAS loci. Across all studies, at least one colocalized eQTL or sQTL was identified in 179 IBD loci (55.9%), implicating 503 unique genes (Supplementary Fig. 6; Supplementary Table 14). Interestingly, 17 of these were sQTL-only loci, emphasizing the importance of considering potential genetic effects on splicing. Overall, these findings indicate that bulk colon tissue analyses contribute to understanding the genetics of IBD.

### GWAS colocalized genes have shared and distinct eQTL and sQTL signals

We further investigated the 183 genes with both eQTL and sQTL and for which at least one QTL colocalized with an IBD GWAS locus (Fig. 2d, bottom two rows) by performing within-gene eQTL-sQTL colocalization. Of these, we identified 105 eQTL-sQTL shared genes (PP4*>* 0.5; Fig. 2d, bottom row) and 78 genes that did not support eQTL-sQTL colocalization (PP4*<* 0.5; Fig. 2d, middle row; Supplementary Tables 15–16). To better understand the relationships between eQTL, sQTL, and GWAS signals, we looked more closely at the colocalization results, including junctions supporting all three comparisons (Supplementary Table 17). For 74 of the 78 genes with no eQTL-sQTL colocalization, only one molecular trait colocalized with GWAS (Supplementary Table 18). Our analysis suggested that for 28 genes (37.8%), there were distinct genetic associations between eQTL and sQTL, and between the non-GWAS QTL and the GWAS signal (posterior probability of distinct genetic associations (PP3)*>* 0.5, coloc-ABF and coloc-SuSiE; Supplementary Tables 19–20). For another 42 genes (56.8%), the colocalization analysis did not identify a genetic association signal in at least one trait (PP1 or PP2 *>* 0.5, coloc-ABF) in at least one of the non-colocalizating comparisons. In the remaining 4 genes (5.4 %), the colocalization analysis was inconclusive (PP3*>* 0.5, coloc.ABF, unresolved, coloc-SuSiE; Supplementary Results). Of the 105 eQTL-sQTL shared genes, 54 showed colocalizaton with GWAS for both the eQTL and sQTL. For 53 of these, there was at least one splice junction that colocalized with both the GWAS locus and the GWAS eGene (Supplementary Table 17).

There were 55 genes where colocalization was detected for two comparisons, but not the third, with 51 showing eQTL-sQTL colocalization (Supplementary Table 21). For 47 genes (85.5%), the third comparison indicated the presence of distinct genetic signals (PP3*>* 0.5, coloc-SuSiE; Supplementary Tables 22–23). Of the remaining eight genes, two (3.6%) reported no genetic association evidence for one of the traits (PP1 or PP2 *>* 0.5, coloc-ABF) and six (10.9%) could not be fine-mapped (unresolved, coloc-SuSiE; Supplementary Table 24). Although we classified colocalization using a PP4 *>* 0.5 threshold, the underlying posterior support is continuous, and differences in association strength, limited fine-mapping resolution, or equivocal support for shared versus distinct signals may affect pairwise classifications and contribute to these 55 genes (Supplementary Results). However, colocalization in two comparisons but not the third may also reflect differences in the splice junctions or association components involved. To illustrate this possibility, we examined the underlying junctions and components, and focused on 42 GWAS sQTL-only genes with eQTL-sQTL colocalization (GWAS-eQTL 39 genes PP3 *>* 0.5, coloc-SuSiE, 3 genes unresolved). Within this group, we identified two scenarios involving different junctions or association components, illustrating the complexity of the genetic regulation. We first found that for 8 genes, colocalization of sQTL with eQTL and separately with GWAS signals involved different splice junctions (Supplementary Table 25). We further found that for 5 genes, there were two distinct sQTLs for the same splice junction, one that colocalized with the eQTL and one with the GWAS signal. For example, at *HAVCR1*, one sQTL colocalized with GWAS (PP4=0.983) while another sQTL for the same junction colocalized with the eQTL (PP4=0.918). Supporting this, colocalization of the GWAS signal with the eQTL showed distinct signals (PP3=0.995). In addition, the candidate shared variants for the two comparisons, rs1363232 (GWAS-sQTL) and rs9790967 (eQTL-sQTL), are in low LD (*r*^2^ *≈* 0; All Populations ^20^). A similar situation could also be found if multiple components are present in the eQTL signal (Supplementary Fig. 7) or GWAS signal (Supplementary Fig. 8).

Together, these analyses provide a glimpse into the complexity of shared and distinct genetic regulation of splicing and expression of the same gene, and their relationship with IBD risk. We acknowledge that in some cases, fine mapping resolution (Supplementary Fig. 9; Supplementary Table 26) or uncertainty in evidence supporting distinct versus shared signals may affect our ability to accurately identify the colocalization status. Considering the junctions, components, and posterior evidence together could clarify which molecular signals share the IBD association and how they relate to the other QTL, while recognizing relationships that remain unresolved and need additional data.

### sQTL variants may lead to different splicing consequences among GWAS colocalized sGenes

Differential splicing can affect RNA abundance, protein abundance, and protein function in multiple distinct ways. Three of these non-mutually exclusive mechanisms are NMD, 5*^′^* UTR remodeling, and protein domain alteration. Biologically, NMD couples alternative splicing to RNA degradation. NMD can occur when an mRNA from a protein-coding gene contains a premature stop codon that a ribosome encounters during translation. This can reduce the abundance of the affected transcript and may lower the total mRNA abundance measured for that gene by RNA-seq. 5*^′^* UTR remodeling may affect translation efficiency through changes in RNA structure or upstream open reading frames (uORFs) that influence translation initiation^21,22^. Proteins often contain subsequences of amino acids (domains) that function together, such as to bind DNA or another protein or to perform a metabolic function. Thus, how sQTL affect protein function depends on which splicing mechanism the target splice junction alters.

To identify potential NMD, Leafcutter2 predicts whether the usage of particular splice junctions leads to non-coding transcripts, which are labeled as an unproductive (UP) splicing event. We classified sQTL as having an NMD-related event when it was predicted to increase or decrease the production of an UP, NMD-sensitive transcript, including events involving exon skipping or poison exon inclusion that introduces a premature stop codon. We defined 5*^′^* UTR remodeling when all junctions in an associated splice cluster were located within an annotated 5*^′^* UTR. Lastly, a protein domain alteration was defined when the inclusion or skipping of an exon overlapped a Pfam annotated protein domain (Supplementary Methods; Fig. 3a). Among the 164 GWAS sGenes with GWAS-sQTL colocalization, 11 (6.7%) were predicted to have an NMD-related event, 55 (33.5%) to have a 5*^′^* UTR remodeling event, and 94 (57.3%) to have a protein domain altering event (Fig. 3b; Supplementary Table 27). Multiple categories could be associated with a single gene because many genes contain multiple splice clusters mapped to a sQTL. We wanted to know which splicing mechanism could be associated with disease. To do this, we tested whether the frequency of these functional mechanisms was enriched within GWAS colocalizing sharing status (Supplementary Fig. 10).

**Figure 3:**
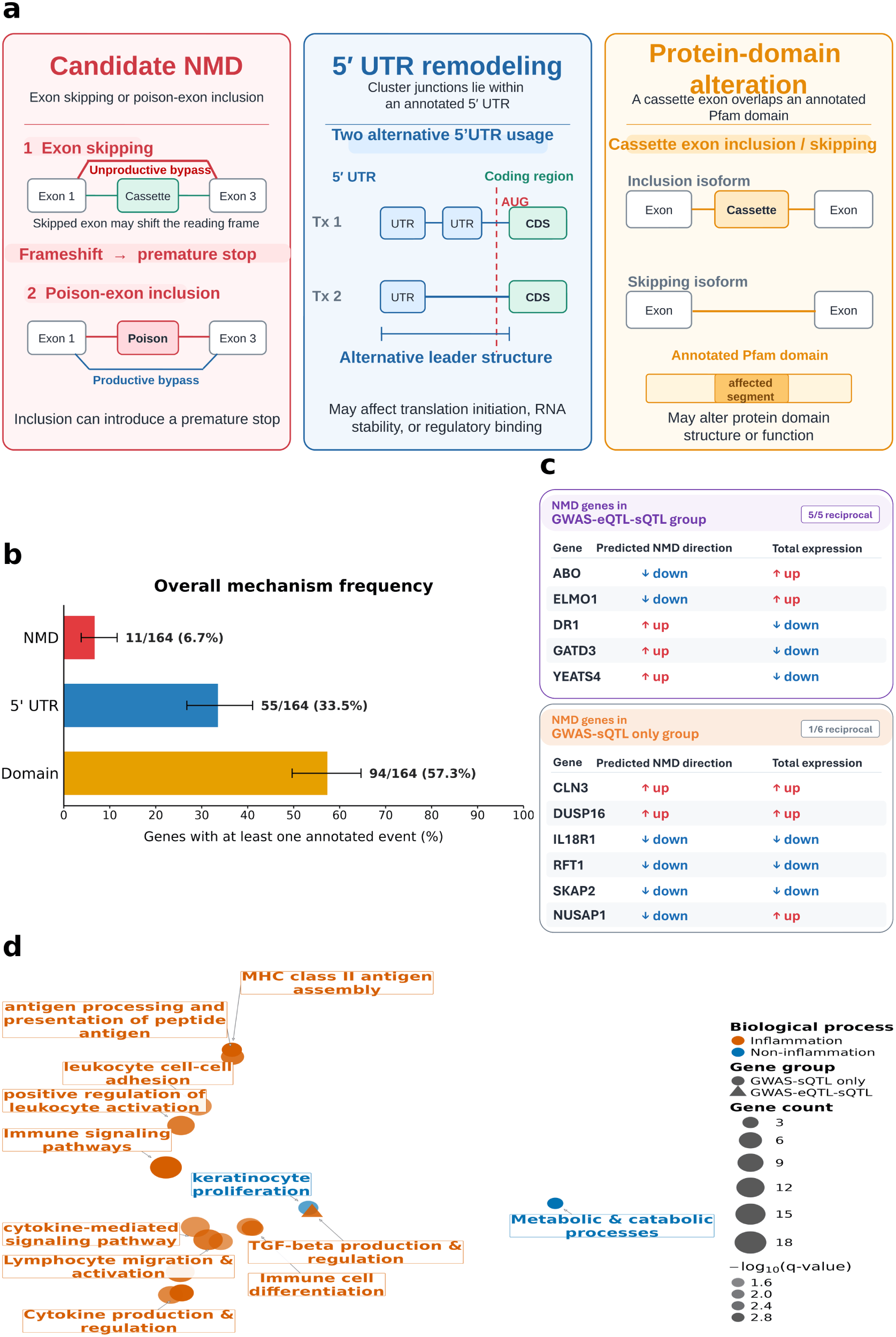
The potential splicing effects of the sGenes that have been colocalized in the GWAS study. **(a)** Illustrative definitions of three candidate splice consequences: nonsense mediated decay (NMD)-sensitive events from exon skipping or poison-exon inclusion, 5*^′^* untranslated region (UTR) remodeling, and exon changes overlapping annotated protein domains. **(b)** Frequencies of the three possible splicing consequences among the 164 sGenes that were colocalized in GWAS studies. **(c)** Direction of NMD-sensitive junction usage relative to total gene expression among the 11 candidate NMD genes across GWAS-eQTL-sQTL group and GWAS-sQTL only group. **(d)** Enrichment of Gene Ontology (GO) biological processes in the disease prioritized sGenes. Dots labeled by enriched processes, where dot size indicates the number of QTL genes in the biological process set and color intensity corresponds to *−* log_10_(*q*-value).

All 11 genes with candidate NMD events had detectable eQTLs, of which 5 had colocalized eQTL and sQTL, and eQTL and GWAS signals and 6 were GWAS-sQTL-only genes. We determined whether gene expression changed in a reciprocal direction as NMD events. Although the frequency of candidate NMD events did not significantly differ across the GWAS-QTL sharing status (all Fisher’s exact *P ≥* 0.39), reciprocal changes in NMD-sensitive junction usage and total expression were observed more frequently in GWAS–eQTL–sQTL genes than in GWAS–sQTL only genes (5/5 versus 1/6). For *ABO*, *ELMO1*, and *NUSAP1* the IBD risk allele was associated with decreased predicted NMD isoforms and increased total gene expression. For *GATD3*, *DR1*, and *YEATS4* the risk allele was associated with increased NMD and decreased total expression. In the remaining five GWAS-sQTL-only genes, junction usage and total expression changed in the same direction (Fig. 3c). Four of these genes (*CLN3*, *IL18R1*, *RFT1*, and *DUSP16*) showed strong support for distinct GWAS and eQTL signals (ABF PP3 *>* 0.91 and SuSiE PP3 *>* 0.99), while this was inconclusive for *SKAP2* (ABF PP3=0.38 and PP4=0.49, with SuSiE unresolved) and with modest support for *NUSAP1* (SuSiE PP3 0.5-0.7). Thus, the combination of colocalization and reciprocal effects could provide stronger support for the shared genetic variants associated with NMD related splicing with changes in total gene expression. In contrast, for gene *NUSAP1*, the direction of the expression association was consistent with NMD, but it remains unclear whether the risk allele associated splicing change accounts for the lower total expression that both share with IBD risk.

Compared with predicted NMD, 5*^′^* UTR remodeling and protein domain associated splicing were observed in a larger proportion of genes, including those without detected eQTLs. Their frequencies did not differ significantly across GWAS-colocalization sharing groups. Unlike the reciprocal relationship observed between predicted NMD and gene expression, these mechanisms showed no consistent relationship with total transcript abundance. For example, among 16 genes with 5*^′^* UTR splicing events predicted to change uORFs numbers, risk alleles favored isoforms with higher predicted uORF counts in both *ELMO1* and *FUT2*, while gene expression increased in *ELMO1* but decreased in *FUT2* (Supplementary Table 28). Similarly, protein domain associated splicing showed heterogeneous relationships with expression. Risk alleles favored retention of an exon in a PH 12 domain in *ELMO1* accompanied by increased expression, whereas increased inclusion of exons in a TNFR c6 domain in *LTBR* was associated with decreased expression. Interestingly, TNFR c6-overlapping events were also observed in *CD40* and *TNFRSF14*; all three genes had both eQTLs and sQTLs colocalizing with GWAS signals. We also identified sQTLs for splicing that involved exons in peptidase domain families in *XPNPEP3*, *ERAP1*, *ERAP2*, *SENP7*, and *CTSW*. Both eQTLs and sQTLs colocalized with GWAS signals for *ERAP2*, *SENP7*, and *CTSW*, whereas only sQTLs colocalized for *XPNPEP3* and *ERAP1*, despite having detected eQTL signals. These findings suggest that 5*^′^* UTR remodeling and protein domain associated splicing may have effects across a broader range disease associated genes than NMD, potentially influencing translation or protein function in ways not limited to changes in total transcript abundance.

### Splicing events in *PTPN2*, *TLR4*, and *ELMO1* illustrate potential contributions of altered splicing to IBD risk

To examine the potential functional consequences of disease associated splicing under different GWAS, eQTL, and sQTL colocalization patterns, we selected three representative genes and examined how changes in junction usage correspond to specific transcripts and their predicted functions. For *PTPN2* gene, a GWAS signal colocalized with an sQTL without a detectable eQTL, illustrating that genetic variation may contribute to disease risk through altered splicing without a detectable association with total gene expression. For *TLR4*, eQTL and sQTL were detected, but the GWAS signal only colocalized with the sQTL, again suggesting that disease associated splicing can occur without a corresponding effect on expression. For *ELMO1*, both eQTL and sQTL signals colocalized with IBD GWAS and with each other, and splicing analysis showed the risk allele may be associated with decreased NMD. This suggests that disease associated splicing changes may be affecting total observed gene expression.

#### *PTPN2* : altered splicing but not expression may lead to reduced expression of a canonical protein

*PTPN2* encodes a T cell protein tyrosine phosphatase, an intracellular regulator of cytokine and immune receptor signaling. Four protein isoforms have been described: PTPN2.1 (transcript *PTPN2*-201), PTPN2.2 (*PTPN2*-202), PTPN2.3 (*PTPN2*-203), and PTPN2.4 (*PTPN2*-210) (Supplementary Fig. 11). The two well characterized canonical transcript isoforms, *PTPN2*-201 and *PTPN2*-202, share the same catalytic tyrosine phosphatase domain (Y Phosphatase), but with different C-terminal sequences that affect their cellular localization^23^. Compared to *PTPN2*-201 and *PTPN2*-202, *PTPN2-210* has an additional exon, exon 6a, which changes the Y Phosphatase domain sequence, introducing a nuclear export signal. This leads to altered protein localization and inflammatory response while retaining phosphatase activity^24^.

In the IBD Plexus data, LeafCutter2 identified a cluster (clu 97897 -) containing 13 junctions, within which an sQTL was found for the junction chr18:12,817,365:12,818,944 (Fig. 4a,c; Supplementary Fig. 12). This sQTL colocalized with an IBD GWAS signal, while no eQTL was detected for this gene (Fig. 4a). GTEx also reported a *PTPN2* sQTL that colocalized with the IBD GWAS signal but no eQTL, providing external support^24,25^ (Fig. 4b). The junction is close to the canonical tyrosine phosphatase domain; thus, its inclusion status may have an effect on this function of PTPN2. The IBD GWAS index variant was rs547268, the lead sQTL was rs12967678, and rs80262450 marked the colocalized sQTL–GWAS signal (Fig. 4a,b; Supplementary Table 29). The GWAS risk allele G is associated with increased significant junction usage, consistent with the association reported in GTEx^25^. This significant junction is compatible with five transcripts: *PTPN2*-205, *PTPN2*-228, *PTPN2*-229, *PTPN2*-234, and *PTPN2*-237 (Fig. 4e). Among these, *PTPN2*-229 and *PTPN2*-205 are protein coding but *PTPN2*-205 lacks a verified coding region, and the remaining three are annotated as NMD transcripts.

**Figure 4:**
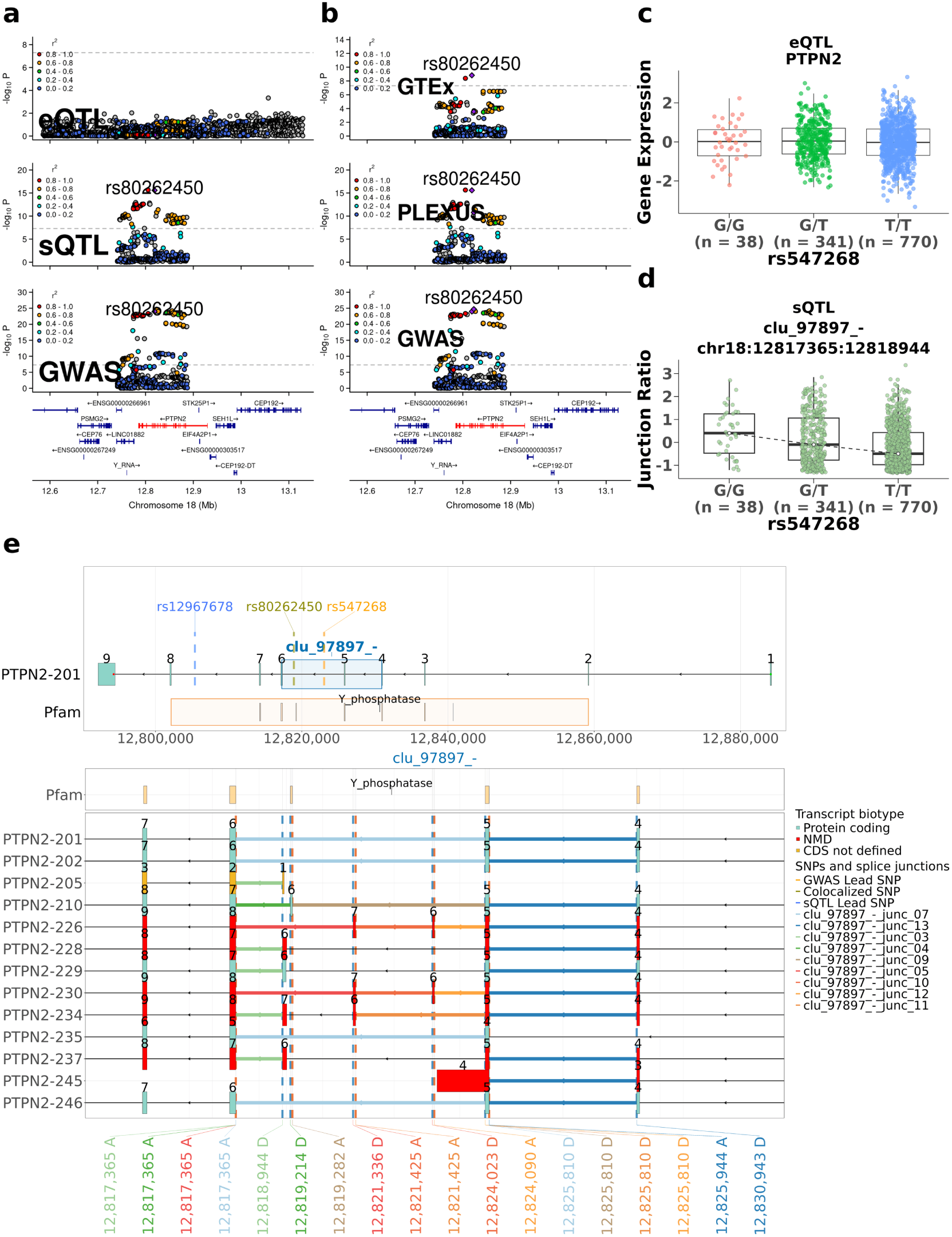
Colocalization of the *PTPN2* sQTL and gene model. **(a)** Locus zoom plots for IBD Plexus *PTPN2* (red gene) eQTL, sQTL and IBD GWAS signals. The purple diamond indicates the shared variant rs80262450 for GWAS-sQTL colocalization; nearby variants are colored according to linkage disequilibrium (*r*^2^) with this variant. The x-axis shows chromosome 18 coordinates, and the y-axis shows *−* log_10_ of the nominal P-values. **(b)** Locus zoom plots for IBD Plexus and GTEx *PTPN2* sQTL and IBD GWAS signals. The purple diamond indicates the shared variant rs80262450 for GWAS-sQTLcolocalization. **(c)** Inverse normalized expression levels of the PTPN2 gene based on genotypes for GWAS index variant rs547268. **(d)** Usage of junction chr18:12817365:12818944 relative to rs547268 genotypes. Panels (c) and (d) display the individual samples superimposed on the box plots, together with the number of samples for each genotype. **(e)**Alternatively spliced GENCODE annotated junctions and transcripts in LeafCutter2-defined cluster clu 97897 - are shown. Intron colors indicate cluster junctions in a transcript. Exon colors indicate the transcript biotype. Vertical dashed lines denote the positions of the GWAS lead variant rs547268 and the colocalized shared variant rs80262450. Positions of Pfam protein tyrosine phosphatase domains indicated at top. **(e)** Overview and enlarged views of the splice cluster in selected GENCODE *PTPN2* transcript models. The upper panel shows the cluster along the reference transcript *PTPN2*-201 and the genomic spans of the Pfam domain, Y phosphatase. Brown, orange, and blue variant markers identify the candidate share variant, GWAS lead variant, and sQTL lead variants, respectively. The lower panels show transcript structures and alternative junctions within three clusters. Exons are colored by transcript annotation as protein-coding, NMD, or CDS not defined. Colored connections identify splice junctions. Exon number in the transcript is displayed above the corresponding exons. Dashed vertical lines mark splice sites in the enlarged views, with genomic coordinates labeled as donor (D) or acceptor (A) sites. Arrows indicate the directionof transcription.

When the *PTPN2*-229 protein coding transcript is expressed, the junction corresponds to inclusion of an additional internal exon relative to the canonical *PTPN2*-202 transcript. This exon introduces an in-frame 41 amino acid (aa) insertion at residues 166–206, producing a 428 aa protein that differs from *PTPN2*-202 within the tyrosine phosphatase domain (Supplementary Table 30). Motif screening identified sequences matching an SH3 binding consensus motif (RHAPLCPDK) and a MAPK docking consensus motif (KRFSHLSL), suggesting the insertion may affect PTPN2 protein interactions. Alternatively, increased usage of the junction may reflect increased expression of one or more of the three NMD-annotated transcripts, potentially affecting transcript stability. The lack of a corresponding eQTL indicating transcript levels were not affected suggests NMD may not play a significant role. Lastly, expression of the non-coding transcript could result in similar gene expression levels being detected, but with reduced production of the protein. Although short read data cannot determine which transcripts result from this splicing event, the GWAS risk allele is associated with increased presence of non-canonical transcripts, potentially increasing non-canonical and reducing canonical protein products.

#### *TLR4* : increased expression of an alternative transcript with altered protein function

*TLR4* encodes the Toll-like receptor type 4 (TLR4) protein whose extracellular leucine-rich repeat (LRR) regions interact with MD-2 to form the LPS (lipopolysaccharide) sensing receptor complex and activate MyD88 and TRIF-dependent inflammatory signaling. The canonical *TLR4*-201 transcript encodes an 839 aa protein. The alternatively spliced *TLR4*-202 transcript includes an additional 5’ UTR exon that results in a downstream shift of the translation initiation site, leading to a 799 aa protein that lacks the first 40 amino acids. This deletion removes the canonical signal peptide and the first 17 residues of the mature extracellular region, including Cys29 and Cys40. Previous studies using N-terminal deletions or substitutions of these cysteines demonstrated impaired MD-2 association, receptor maturation, cell surface expression, and LPS induced signaling^26,27^. Using transcript specific real time PCR, Jarěsová et al.^28^ found reduced LPS-induced upregulation of *TLR4*-202 in cystic fibrosis monocytes and suggested that *TLR4*-202 might encode an inhibitory protein.

In the IBD Plexus data, LeafCutter2 identified a cluster (clu 53716 +) containing five overlapping junctions within *TLR4*, with sQTL associated with two of these, chr9:117,708,318:117,708,563 (Fig. 5c) and chr9:117,704,565:117,708,199 (Supplementary Fig. 13), that shared the same lead variant, rs11536871 (Supplementary Table 31). Only the sQTL for the first junction colocalized with an IBD GWAS signal (PP4=0.97) (Fig. 5a). Although an eQTL was detected for *TLR4*, it colocalized with neither the IBD GWAS signal (PP1=0.63) nor this sQTL (PP2=0.63), with in both cases the eQTL signal not being identified (Fig. 5a,b). The IBD GWAS risk G allele at rs4986790 was associated with increased usage of both junctions, which are present in *TLR4*-202 spanning exons 1-2 and 2-3, respectively (Fig. 5d). The increased usage of this splicing pattern may indicate higher expression of *TLR4*-202, while the lack of a corresponding eQTL suggests this may also lower expression of *TLR4*-201, leading to altered signaling and LPS responsiveness.

**Figure 5:**
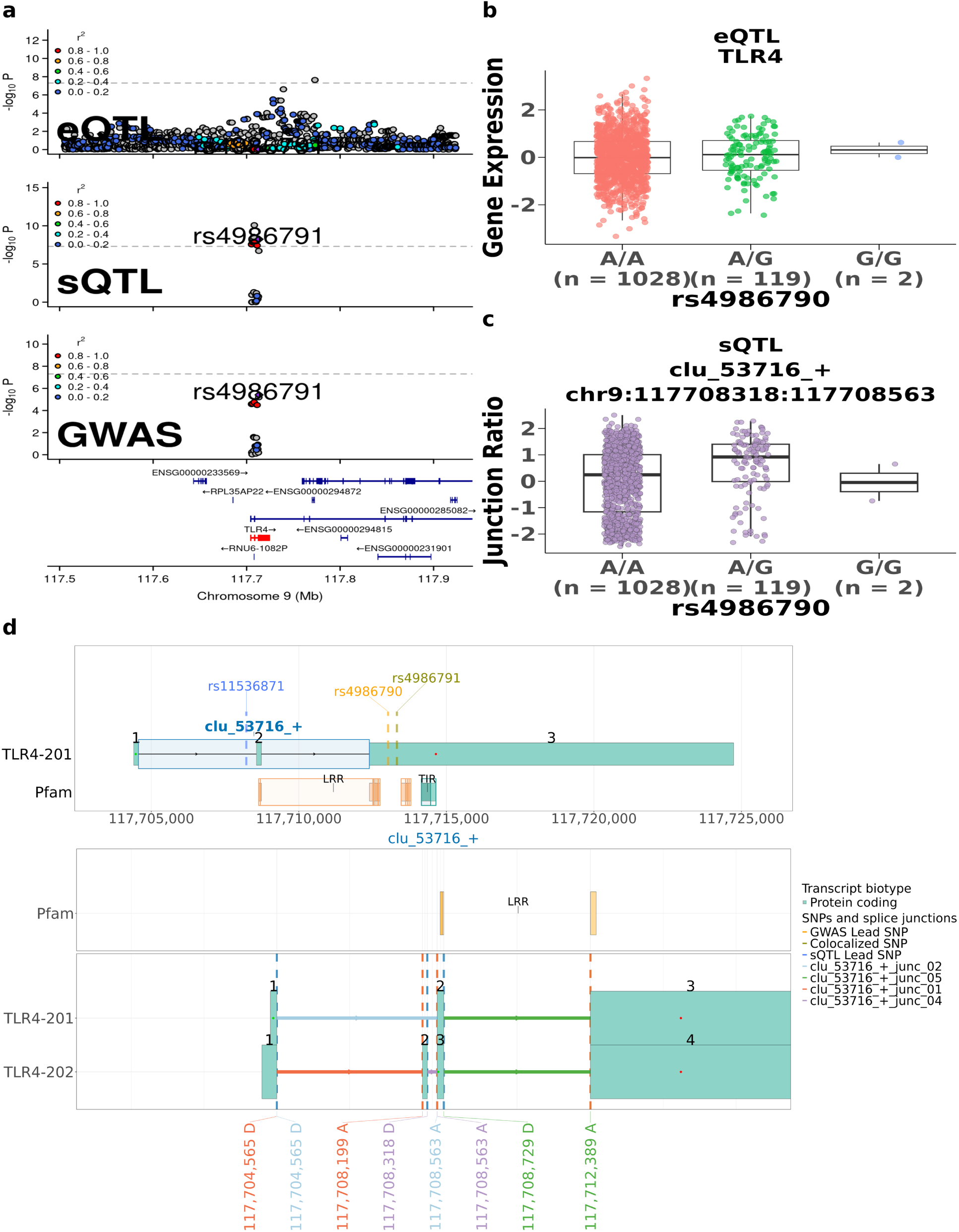
*TLR4* sQTL colocalization and the gene model. **(a)** Locus zoom plots for IBD Plexus *TLR4* (red gene) eQTL, sQTL, and IBD GWAS signals. Purple diamond indicates the GWAS-sQTL colocalization shared variant rs4986791. Nearby variants are colored according to their linkage disequilibrium (*r*^2^) with this variant. The x-axis shows chromosome 9 coordinates, and the y-axis displays *−* log_10_ of the nominal P-values. **(b)** Inverse normalized expression of *TLR4* relative to GWAS lead variant rs4986790 genotypes. **(c)** GWAS-colocalized junction chr9:117708318:117708563 usage in relative to the rs4986790 genotypes. Panels (b) and (c) display the individual samples superimposed on the box plots, together with the number of samples for each genotype. **(d)** Overview and enlarged views of the splice cluster in selected GENCODE *TLR4* transcript models. The upper panel shows the clusters along the reference transcript *TLR4*-201 and the genomic spans of the Pfam domains, LRR and TIR. Brown, orange, and blue variant markers identify the candidate shared variant, GWAS lead variant, and sQTL lead variants, respectively. The lower panels show transcript structures and alternative junctions within the cluster. Colored connections identify splice junctions. Exon numbers in the transcript are displayed above the corresponding exons. Dashed vertical lines mark splice sites in the enlarged views, with genomic coordinates labeled as donor (D) or acceptor (A) sites. Arrows indicate the direction of transcription.

#### *ELMO1* : sQTL may lead to both reduced NMD and 5’ UTR structure affecting protein expression

Engulfment and cell motility protein 1 (ELMO1) interacts with DOCK proteins to activate the Rac1 pathway and has a role in actin remodeling, bacterial engulfment, and immune cell migration^29^. In addition to the canonical transcript *ELMO1*-201, several isoforms exist (Supplementary Fig. 14). These include *ELMO1*-236, which skips exon 18 of the *ELMO1*-201 transcript, causing a frameshift^30^ that potentially introduces a premature stop codon; *ELMO1*-230 that uses an alternative splice site at exon 16; and *ELMO1*-204 that includes an exon overlapping the CED12 encoding region, all three of which are annotated as leading to NMD in GENCODE. Similar to *ELMO1*-236, *ELMO1*-214 also skips exon 18 and is annotated as a non-coding transcript in GENCODE, but for which it has also been suggested that it leads to a protein coding transcript where this skipped exon alters the C-terminal PH-domain coding sequence and thus changes the DOCK/RAC signaling and microbial sensing ^29^. Lastly, the *ELMO1*-225, *ELMO1*-227, *ELMO1*-223 and *ELMO1*-207 transcripts all preserve the canonical coding region seen in *ELMO1*-201 but have alternative 5’ UTR structures. *ELMO1*-223 specifically contains an additional 966 nt 5*^′^* UTR exon with 19 predicted upstream open reading frames (uORFs) compared with two in ELMO1-201, representing a net gain of 17 uORFs, including three additional strong Kozak uORFs (Supplementary Table 28).

In the IBD Plexus data, we identified an eQTL and seven sQTL corresponding to junctions from three splicing clusters that collectively contained 13 junctions, all of which colocalized with the IBD GWAS signal (all sQTL PP4*>* 0.59; eQTL PP4: 0.60). The seven sQTL also colocalized with the eQTL (PP4*>* 0.88) (Supplementary Tables 11–12, 15). One splicing cluster (clu 45751 -) contained an sQTL for a predicted unproductive junction at chr7:36878117:36894854 used in transcripts *ELMO1*-214 and *ELMO1*-236 (Fig. 6a,c,f; variant annotations in Supplementary Table 32). The colocalized GWAS risk G allele at variant rs73120731 was associated with higher total *ELMO1* expression and lower usage of the predicted unproductive junction, indicating reduced expression of *ELMO1*-236 and/or *ELMO1*-214. Three additional junctions were found in splicing cluster (clu 45755 -), of which two, chr7:37013435:37033347, and chr7:37033412:37096619, are compatible with *ELMO1*-204 and a third, chr7:37013431:37096619, with *ELMO1*-230 (Fig. 6d; variant annotations in Supplementary Table 33). The GWAS risk G allele is also associated with decreased usage of all three splice junctions (Supplementary Fig. 15), as expected with predicted NMD for these transcripts. The remaining three junctions were identified in splicing cluster clu 45758 -, of which one, chr7:37342763:37352313 is compatible with *ELMO1*-225 and *ELMO1*- 227, and the other two, chr7:37342763:37353032 and chr7:37342763:37428791, are compatible with *ELMO1*-207 and *ELMO1*-223, respectively (Fig. 6f; variant annotations in Supplementary Table 34). The GWAS risk G allele was associated with increased usage of the junction in *ELMO1*-223 (Fig. 6e), indicating increased transcription of *ELMO1*-223 that contains an increased number of uORFs. Together, these findings illustrate multiple mechanisms by which GWAS-associated splicing changes may alter transcript and protein abundance through NMD and alternative 5*^′^* UTR structures.

**Figure 6:**
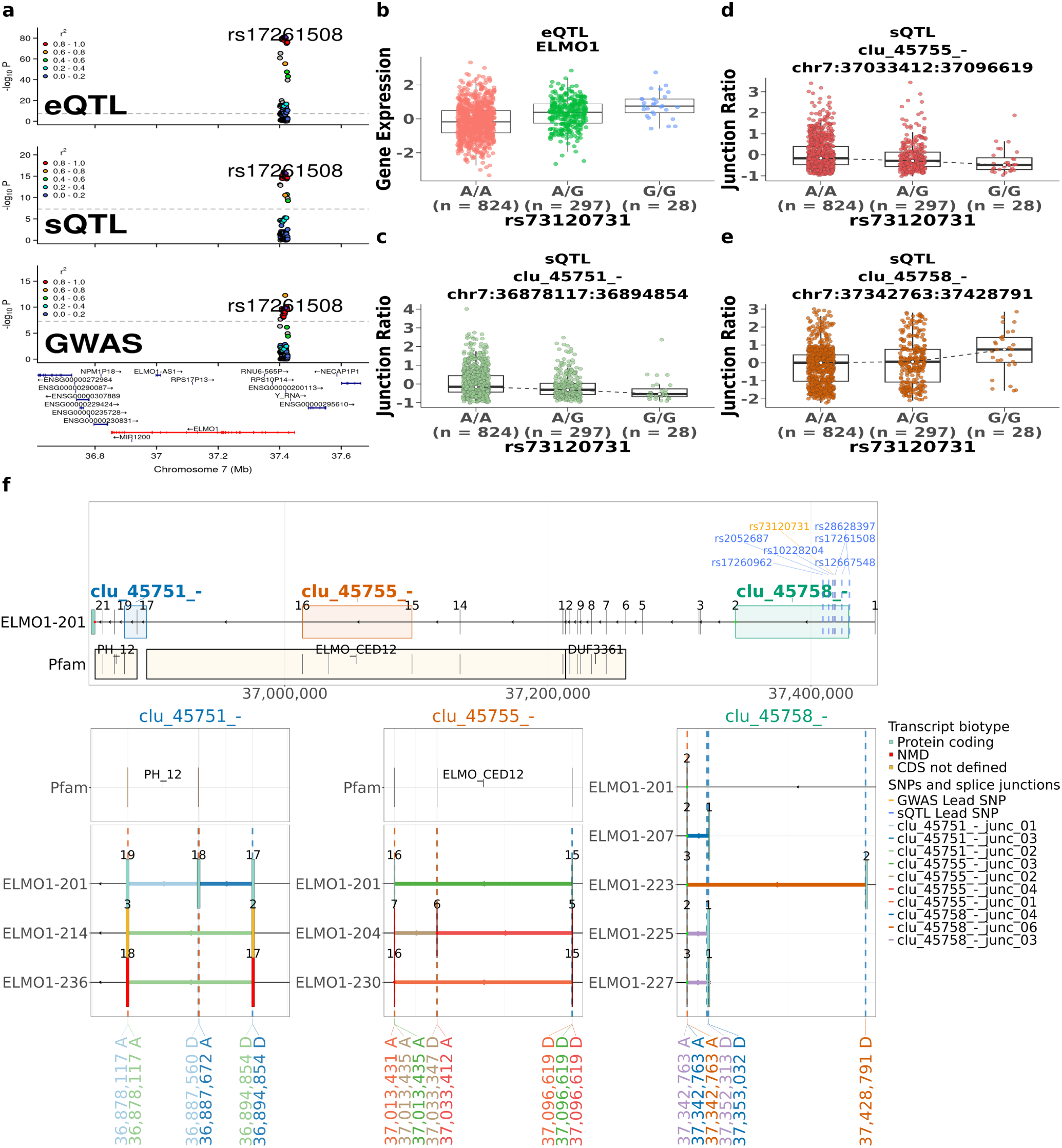
GWAS-colocalized expression and splicing associations at *ELMO1*. **(a)** Locus zoom plots for IBD Plexus *ELMO1* (red gene) eQTL, sQTL, and IBD GWAS signals. A purple diamond indicates the GWAS-sQTL colocalization shared variant rs17261508. Nearby variants are colored according to their linkage disequilibrium (*r*^2^) with this variant. The x-axis gives the chromosome 7 coordinates, and the y-axis displays the *−* log_10_ of the nominal P values. **(b)** Inverse-normalized expression of *ELMO1* relative to GWAS lead variant rs73120731 genotypes. **(c to e)** Usage of three *ELMO1* junctions with sQTL: chr7:36878117:36894854 in cluster clu 45751 - (c) chr7:37033412:37096619 in clu 45755 - (d), and chr7:37342763:37428791 in clu 45758 - (e). Panels (b to e) display individual samples superimposed on the box plots, with the samples stratified by rs73120731 genotype. **(f)** Overview and enlarged views of the three splice clusters in selected GENCODE *ELMO1* transcript models. The upper panel shows the clusters along the reference transcript *ELMO1*-201 and the genomic spans of the Pfam domains PH 12, ELMO CED12, and DUF3361. Orange and blue variant markers identify the GWAS lead variant and sQTL lead variants, respectively. The lower panels show transcript structures and alternative junctions within each cluster, with Pfam tracks above the first two clusters. Exons are colored by transcript annotation as protein coding, NMD, or CDS not defined. Colored connections identify splice junctions. Exon number in transcript displayed above the corresponding exons. Dashed vertical lines mark splice sites in the enlarged views, with genomic coordinates labeled as donor (D) or acceptor (A) sites. Arrows indicate the direction of transcription.

### IBD colocalized sGenes are associated with immune response and are drug targets

To understand cellular processes that may be affected by GWAS colocalized eGenes and sGenes, we first performed Gene Ontology enrichment using a background of all GO-annotated eGenes and sGenes (Fig. 3d). We found that the GWAS-colocalized sGenes were enriched for several immune-related pathways including cytokine production and signaling, adaptive immune responses, dendritic cell differentiation, and immune effector functions (Supplementary Table 35). This may reflect the sensitivity of immune functions to alternative splicing, as changes in even a small protein segment can affect bacterial sensing, receptor–ligand binding, or downstream signaling. In contrast, the GWAS-colocalized eGenes showed no significant pathway enrichment. To alternatively demonstrate the range of potential effects of eGenes and sGenes on IBD risk, we organized them within a descriptive framework focused on biological functions implicated in IBD pathogenesis (Supplementary Results; Supplementary Table 36), including mucosal and microbial interactions, epithelial responses, microbial and antigen sensing, intracellular inflammatory signaling, and immune cell activation and trafficking^31–33^. These show that genetic predisposition for IBD is linked to a heterogeneous mix of immune and epithelial cell processes previously associated with disease.

Many existing IBD drugs target immune related proteins, and efficacy of these therapies can vary widely across patients. Given the enrichment of sGenes for immune related pathways, we wondered whether our analyses showed evidence of sQTL that may influence splicing in targeted proteins. *CD40*, which has both eQTL and sQTL that colocalize with a GWAS signal, encodes the target of anti-CD40 antibodies that have been evaluated in IBD. The canonical full-length transcript, *CD40*-202, contains 4 cysteine-rich domains (CRDs) (Supplementary Fig. 16). We found that the GWAS risk T allele for variant rs1883832 was associated with decreased *CD40* expression and increased usage of a splice junction that was compatible with two alternative transcripts, *CD40*-217 and *CD40*-223, that skip exons encoding CRD3 and 4, while retaining CRD1 and 2 (Supplementary Fig. 17; Supplementary Table 37 for variant annotations). Anti-CD40 antibodies have been designed to recognize different extracellular CRDs, including Lob 7/6 that targets CRD3^34^, ravagalimab that targets CRD2^35^, and ch5D12 that targets CRD1^36^. Therefore, increased expression of transcripts lacking CRD3 and CRD4 in patients with the risk allele may affect response to antibodies targeting these two domains, suggesting patient genotypes could inform drug use.

To place the proposed IBD risk genes in a therapeutic context, we used drug–target and indication annotations from the Open Targets Platform to identify genes targeted by drugs investigated or approved for IBD or for other diseases (Fig. 7). We also reviewed published studies and clinical trial registrations to identify drugs investigated for UC or CD whose IBD indications were not represented in the Open Targets data used here (release 26.09), including drugs targeting *CD40*, *ITGAL*, and *IL22RA1* (Fig. 7; Supplementary Table 38). We then examined how disease associated splicing might affect protein features relevant to drug action. As described above, the canonical *TLR4*-201 transcript encodes a protein that interacts with MD-2 to form a LPS sensing receptor complex, but for which we predict a sQTL colocalized with the GWAS risk allele leads to increased expression of *TLR4*-202, encoding a shorter protein that impairs MD-2 association, potentially reducing the amount of functional TLR4–MD–2 complex^27^. The drug eritoran inhibits TLR4 signaling by binding MD-2, whereas resatorvid (TAK-242) targets an intra-cellular domain of TLR4^37,38^. Splicing associated genotype variation in drug response has a precedent in therapies that target the gene *HMGCR* ^39^. Our observations raise a similar possibility for *TLR4*; genotype associated differences in transcript usage could influence responses to drugs acting at different sites within the TLR4–MD-2 signaling complex, although this hypothesis remains to be tested.

**Figure 7:**
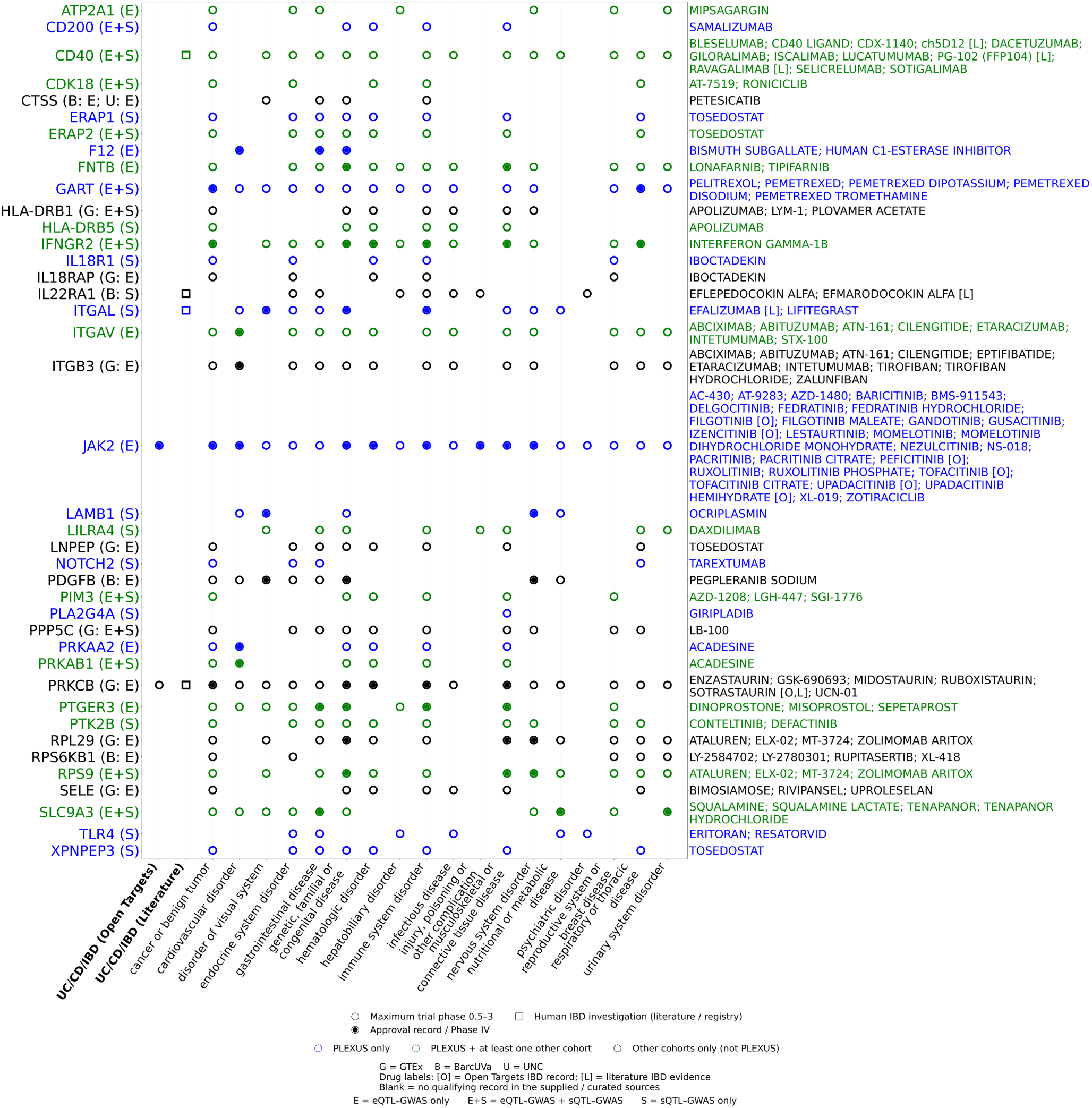
Comparison between the disease effector genes identified by colocalization in IBD Plexus, UNC, GTEx, and BarcUVa. The maximum stage of clinical trials therapeutic modifiers of each gene are described to be at in the Open Target Genes (Methods). Hollow circles indicate a maximum recorded trial phase of 0.5–3 without an approval or phase-IV record in the corresponding gene–disease-group cell. Solid circles indicate that at least one contributing drug has an approval record or phase-IV evidence within that disease group. Multiple drugs may contribute to each cell. The UC/CD/IBD (Open Targets) and UC/CD/IBD (Literature) columns indicate IBD investigations or approvals recorded in Open Targets and human IBD investigations documented in published studies or clinical trial registrations, respectively; hollow squares denote the latter. Supporting drugs are individually labeled [O] for Open Targets, [L] for literature or trial registry evidence, or [O, L] for both; these annotations apply to the marked drugs rather than every drug listed in the corresponding gene row. Blue indicates genes nominated only in IBD Plexus, green indicates genes nominated in IBD Plexus and at least one other cohort, and black indicates genes nominated in other cohorts but not IBD Plexus. Gene-label annotations indicate eQTL–GWAS colocalization only (E), sQTL–GWAS colocalization only (S), or both (E+S).

## Discussion

In a recent study, genetic heritability was estimated to be 78% for Crohn’s disease and 57% for ulcerative colitis, most of which has not been explained in detail, emphasizing the ongoing need to understand how genetic variation contributes to IBD ^40^. In this study, we showed that an increased cohort of 1,149 IBD colon samples and a focus on both expression and splicing were able to substantially increase the number of potential disease genes and molecular mechanisms underlying IBD risk. Specifically, we prioritized 345 genes, including 164 sGenes and 239 eGenes across 147 IBD GWAS loci, nearly half of the 320 IBD GWAS loci (45.9%). Together with previous bulk colon cohorts, colocalized molecular QTLs were identified at 179 loci (55.9%), implicating 503 genes. These findings suggest that as QTL studies become larger, include more diverse patient populations, and include more molecular traits, we will increasingly be able to predict molecular mechanisms underlying most IBD GWAS loci.

Our analyses also support a growing appreciation for the complexity of how genetic variants affect IBD risk. Over half of IBD GWAS loci that we colocalized with QTLs involved more than one gene, suggesting many risk variants, alone or in combination with variants in high linkage disequilibrium, affect expression and splicing of multiple genes in a single locus. In addition, as many predicted target genes were non-coding and could themselves affect expression of nearby genes, variants may indirectly affect regulation of disease genes. Although more than one target gene in a single locus may contribute independently or jointly to IBD risk, in other cases this may simply reflect shared genetic regulation of non-disease genes. Experiments that capture altered regulatory effects on both individual genes and in conjunction with others within the same locus will be needed to understand the effects of variants. Therefore, a comprehensive interpretation of IBD risk loci must consider all genes supported by molecular evidence and not focus on a single causal target.

There is a growing appreciation for the role of alternative splicing in disease. For many genes, their role in cellular functions is not simply based on their overall level of expression, but more specifically on the expression of particular isoforms that can alter the production and function of canonical protein products. In this study, we showed that sQTLs may affect the frequency of at least three potential splicing-related mechanisms, namely NMD, 5*^′^* UTR remodeling, and protein domain alteration. It is also important to note that multiple splicing mechanisms can be affected by genetic variation within the same gene. As shown by *ELMO1*, different junctions associated with sQTLs could involve potential NMD, 5*^′^* UTR remodeling, and protein domain alteration. Each of these can affect transcription in different ways, namely transcript stability, translation efficiency, and protein function, respectively. These emphasize that investigations of the roles of particular genes in disease must consider more than just total expression, and that effects of alternative splicing may need to be tested in multiple ways.

While GWAS colocalized eGenes represented a range of cellular functions, we found that sGenes were enriched for immune-related processes, suggesting a potential role for alternative splicing in immune regulation. These findings may also have implications for drugs targeting immune-related proteins, as alternative splicing can alter protein domains involved in drug binding or signaling. Our analysis of *CD40* and *TLR4* raises the possibility that genotype associated differences in transcript usage could affect these interactions. A precedent has been reported for the gene *HMGCR*, where rs3846662 was associated with alternative splicing of exon 13, which encodes part of statin-binding domain. A greater induction of the exon-skipped transcript in patient derived cells was associated with smaller LDL-cholesterol reduction during simvastatin treatment^39^. In prostate cancer, detection of the androgen-receptor splice variant AR-V7, which lacks the ligand binding domain, was associated with resistance to enzalutamide and abiraterone ^41^.

Although the latter does not establish a genotype dose effect, both studies illustrate how differences in target isoforms may be relevant to treatment response. Whether similar relationships exist for the IBD associated splicing events identified here remains to be tested. Functional studies and analyses integrating genotype, altering protein domains, and treatment outcomes would help determine whether these splicing differences have implications for drug response before considering their use in treatment selection or trial recruitment.

We acknowledge several limitations to this study that should be considered. First, although our study included 1,149 IBD colon samples, the sample size remains limited for more detailed analyses of QTL effects across different IBD phenotypes, such as disease activity, inflammatory status, and clinical subgroups. Larger cohorts will be needed to determine whether these genetic effects differ across disease phenotypes. Second, transcription data was derived from bulk colon tissue. While we attempted to reduce the effects of technical and biological covariates reflecting differences in cell composition and inflammatory state that may influence observed gene expression and splicing patterns, these still may have impacted our results. In addition, these data cannot determine whether observed effects are specific to individual cell types. Cell type specific QTL mapping will better determine which cells are responsible for these signals. Third, using short read RNA-seq, we concentrated on the usage of splice junctions, not the full length transcripts. Long-read sequencing will be important for identifying more specifically which isoforms are affected by the sQTLs. Fourth, the splicing mechanisms and their associated consequences were predicted based on known transcripts annotated by GENCODE. Unannotated transcripts may contribute to the observed junction usage changes with different functional consequences that could not be predicted here. Lastly, colocalization identifies shared genetic signals but cannot prove that changes in expression or splicing are related to IBD risk. Targeted genetic studies are necessary to show this more definitively.

## Conclusion

In summary, combined eQTL and sQTL mapping in 1,149 colon tissue samples from IBD patients revealed evidence of genetic effects on expression and/or splicing associated with almost half of the tested IBD GWAS loci. Importantly, a more in depth analysis of splicing effects showed the complex genetic influences on disease, including by influencing RNA abundance, translation efficiency, and protein function beyond changes in total gene expression. Splicing changes might more specifically affect immune signaling and could potentially impact drug response. Large and more diverse patient cohorts, along with long read sequencing and functional studies, will be needed to better understand these mechanisms and their possible application in the treatment of IBD.

## Methods

### Study population and molecular data

We obtained existing 150-bp short read RNA Sequencing (RNA-seq) data and genotype data from the Crohn’s & Colitis Foundation IBD Plexus platform ^42^. The initial dataset comprised 1,197 colonic samples from patients with ulcerative colitis (UC), Crohn’s disease (CD), or IBD diagnosis. Genotype data were generated using the Infinium Global Screening Array and imputed with the TopMed-r2 reference panel. Variants with an imputation quality score of *r*^2^ *<* 0.3 or an allele frequency below 0.01 were excluded. RNA-seq reads were aligned to the GRCh38 human reference genome using STAR with the GENCODE v49 gene annotation. The WASP option for STAR was used to reduce allelic mapping bias during read processing. VerifyBAMID was used to assess sample identity of the RNA-seq, and samples with identity metrics *<* 0.8 were excluded. Following these quality control measures, 1,149 samples remained in the final analytic dataset.

### Molecular trait quantification

STAR alignment output files were used to quantify gene expression (bam) and splice junction usage (SJ.out.tab). For splice junction usage, we used LeafCutter2, which grouped splice junctions sharing donor or acceptor sites into clusters. Within each cluster, junction usage was calculated as the percent spliced in (PSI), the ratio of reads supporting a junction to the total number of reads supporting all junctions in that cluster. LeafCutter2 annotated each junction as productive (PR), unproductive (UP), neither productive nor unproductive, or intergenic. Junctions and clusters were assigned to genes using GENCODE v49 transcript models. Gene expression was quantified using QTLtools, and junctions were required to have at least 6 reads in *>* 20% of samples, converted to counts per million, and adjusted by the trimmed mean of M values. Outlier genes in the expression matrix and clusters in the PSI matrix identified by eQTLQC were removed.

### eQTL and sQTL mapping and fine mapping

Gene expression values were inverse-normal transformed, and PSI values were quantile-normalized. We used QTLtools to test variants within 1 Mb of each gene transcription start site for eQTL mapping and within 1 Mb of each splice junction for sQTL mapping. Models included collection year, the top four genotype principal components, and unwanted hidden factors estimated with RUVSeq. The final models included 26 RUVSeq factors for gene expression and three for PSI. QTLtools permutation analyses used 1,000 permutations, and empirical thresholds were estimated with the Storey and Tibshirani method^43^. Primary discovery results were reported at 5% FDR. Genes passing the eQTL threshold were called eGenes, significant junctions were called sJuncs, and genes with at least one sJunc were called sGenes.For each eGene and sJunc, the associated variant with the smallest nominal *P* value was selected as the lead QTL variant. A gene level lead sQTL was the strongest variant–junction association assigned to the sGene. QTLtools conditional analysis separated primary and secondary signals. SuSiE was used to fine map significant eGenes and sJuncs and to generate 95% credible sets using LD calculated from IBD Plexus genotypes^44^.

### Credible set sharing and genomic characteristics

Credible set variant-gene pairs were classified as eQTL-only, sQTL-only, or shared. Unique variant analyses additionally distinguished mixed QTL variants associated with expression and splicing of different genes. Groups were compared for TSS and splice site distances and absolute QTL effects. Deduplication and classification rules are detailed in Supplementary Methods.

### Cross Cohort QTL Comparison

We compared IBD Plexus molecular QTLs with colon eQTL and sQTL data from GTEx and BarcUVa and with diseased-colon eQTL data from UNC^7,11,12^. Replication was assessed using exact variant–gene or variant–junction matches and LD proxies with *r*^2^ *≥* 0.8. Effect size concordance was evaluated for matched associations. TopLD was used to identify LD proxies in African (AFR), East Asian (EAS), European (EUR), and South Asian (SAS) reference populations^45^.

### Molecular QTL and IBD GWAS colocalization

We used a two stage hybrid colocalization framework implemented in the coloc R package. Each pair of traits was first analyzed using coloc.abf, which assumes at most one causal signal per trait within the tested region. Pairs not considered colocalized (PP4*<* 0.5) were subsequently evaluated with coloc.susie when a matched LD matrix was available. A trait pair was considered colocalized when PP4 exceeded the threshold (*>* 0.5) in coloc.abf or in at least one coloc.susie signal pair.

We obtained IBD GWAS summary statistics and lead variants for 320 loci^4^. A molecular QTL and GWAS signal was selected when the lead eQTL or lead sQTL was in LD with the GWAS lead variant at *r*^2^ *≥* 0.2 in at least one TopLD reference population or when a GWAS variant was present among the molecular QTL variants. Summary statistics were restricted to the GWAS defined LD block, and the hybrid colocalization framework was applied. A disease prioritized eGene or sGene had at least one eQTL–GWAS or junction level sQTL–GWAS comparison with PP4*>* 0.5, respectively. We applied the same pairing criteria and hybrid colocalization framework to IBD Plexus, GTEx, BarcUVa, and UNC. We additionally performed cross-cohort rescue testing: when an eQTL–GWAS or sQTL–GWAS pair colocalized in any cohort, we tested the corresponding pair in the remaining cohorts using the same framework when the required summary statistics were available.

### Gene groups and component-level comparisons

Genes were grouped by GWAS colocalization with expression, splicing, or both, and by within-gene eQTL–sQTL sharing. Non-colocalizing comparisons were classified using ABF/SuSiE evidence for distinct, single-trait, or inconclusive associations. Junctions, components, candidate shared variants, and LD were compared across traits. Locus multiplicity, gene biotypes, and cohort overlap were summarized using rules detailed in Supplementary Methods.

### Functional annotation, enrichment

Variants were annotated with snpEff, Ensembl VEP, MaxEntScan, SpliceAI, PolyPhen, and SIFT. We tested functional enrichment using non-QTL background variants matched by minor allele frequency and distance to the transcription start site. Regulatory element analyses included eCLIP peaks for 55 splicing related RNA binding proteins, consensus histone mark regions, and Pfam protein domain annotation. Matched null regions were generated with bootRanges, and multiple testing was controlled across enrichment analyses. Gene Ontology biological process enrichment was performed for GWAS colocalized genes using all detected QTL genes as the background.

### Candidate splicing mechanisms and interpretation

We tested splicing mechanisms only on GWAS colocalized junctions. Candidate NMD events were identified from LeafCutter2 UP annotations and transcript level premature termination codon rules. We additionally annotated clusters occurring within 5*^′^* UTRs and exon skipping events overlapping Pfam protein domains. For NMD candidates, expression and junction usage effects were aligned to the same IBD risk allele and classified as reciprocal or same direction. For 5*^′^* UTR events, we further evaluated uORF prediction and the Kozak-context classification. Candidate locus interpretation used GENCODE v50 transcript models, Pfam domain, genotype stratified expression, and PSI values, and transcript/CDS comparisons. Analyses included uORF prediction, Kozak context classification, and protein motif screening.

### Mapping disease prioritized genes to drug targets

We mapped genes with eQTL or sQTL signals colocalized with IBD GWAS signals (PP4 *>* 0.5) to drugs in the Open Targets Platform. Gene–drug pairs were retained based on mechanism of action target annotations and qualifying clinical development or approval records. We excluded IBD related indications, selected non-disease categories, and four known targeted genes (*JAK2*, *PRKCB*, *CD40*, and *ITGAL*) for IBD treatments from the final visualization.

### Data visualization and statistical analysis

We generated figures in R and Python. ggtranscript generated transcript model plots. locusZoomR (v0.2.0) generated regional colocalization plots. matplotlib generated the drug targeted figure. We used Fisher’s exact tests to calculate odds ratios for categorical enrichment analyses. We adjusted multiple tests with the Benjamini-Hochberg method. Unless stated otherwise, all statistical tests were two-sided.

## Data availability

IBD Plexus data are available upon approved application to Crohn’s & Colitis Foundation IBD Plexus program (https://www.crohnscolitisfoundation.org/ibd-plexus). Publicly available v8 eQTL and sQTL summary statistics were downloaded from the GTEx portal (https://gtexportal.org/home/downloads/adultgtex/overview). Full summary statistics for BarcUVa-seq eQTL and sQTL were downloaded from the Digital Repository of the University of Barcelona (https://diposit.ub.edu/dspace/handle/2445/172697). Source data are provided with this paper. Lead summary statistics and associated results are provided in Supplementary Data 1 (worksheets S1–S44) and the Source Data file.

## Web Resources

Crohn’s & Colitis Foundation IBD Plexus, https://www.crohnscolitisfoundation.org/research/plexus.

NHLBI Trans-Omics for Precision Medicine (TOPMed) Program, https://topmed.nhlbi.nih.gov/.

TOPMed Imputation Server, https://imputation.biodatacatalyst.nhlbi.nih.gov/.

NCBI Database of Single Nucleotide Polymorphisms (dbSNP), https://www.ncbi.nlm.nih.gov/snp/.

GENCODE Human Release 49, https://www.gencodegenes.org/human/release_49.html.

GTEx Portal, Adult GTEx data downloads, https://gtexportal.org/home/downloads/adult-gtex/overview.

BarcUVa-Seq eQTL and sQTL summary statistics, https://hdl.handle.net/2445/172697.

TOP-LD web server, http://topld.genetics.unc.edu/.

TOP-LD API documentation, https://github.com/linnabrown/topld_api.

UK Biobank, https://www.ukbiobank.ac.uk/use-our-data/.

Open Targets Platform, https://platform.opentargets.org/.

Ensembl Variant Effect Predictor (VEP), https://www.ensembl.org/info/docs/tools/vep/index.html.

ENCODE Portal, https://www.encodeproject.org/.

UCSC Genome Browser, https://genome.ucsc.edu/.

Gene Ontology Resource, https://www.geneontology.org/.

Pfam protein-family and protein-domain annotations through InterPro, https://www.ebi.ac.uk/interpro/ entry/pfam/.

IBD Plexus eQTL–sQTL analysis code, https://github.com/REPLACE_ACCOUNT/REPLACE_REPOSITORY.

## Code Availability

eQTL and sQTL pipelines are available on GitHub https://github.com/yuchen-hahaha/sQTL-pipeline as well as Code Ocean.

## Author contributions

E.Y.-C.H., T.S.F., and S.Z.S. conceptualized study. E.Y.-C.H., S.H., N.H.H., and N.C.N contributed to data pre-processing. E.Y.-C.H. contributed to data analysis. E.Y.-C.H., S.Z.S, and T.S.F. prepared the manuscript. All authors reviewed the manuscript.

## Acknowledgements

The authors thank the participants, clinicians, and investigators of the Study of a Prospective Adult Research Cohort with IBD (SPARC IBD) who have contributed time, data, and samples. SPARC IBD is a part of the Crohn’s & Colitis Foundation’s IBD Plexus data sharing platform that includes data from electronic health records and biospecimens. This study was supported by the National Institute of Diabetes and Digestive and Kidney Diseases of the National Institutes of Health (NIH) Award Numbers F31DK137574, 1R01DK136262, 1R01DK138462.

