## Supplemental Information for "Beyond Abundance: Integrated Splicing and Expression Quantitative Trait Loci Refine Causal Gene Mapping in IBD"

### Supplementary Figures

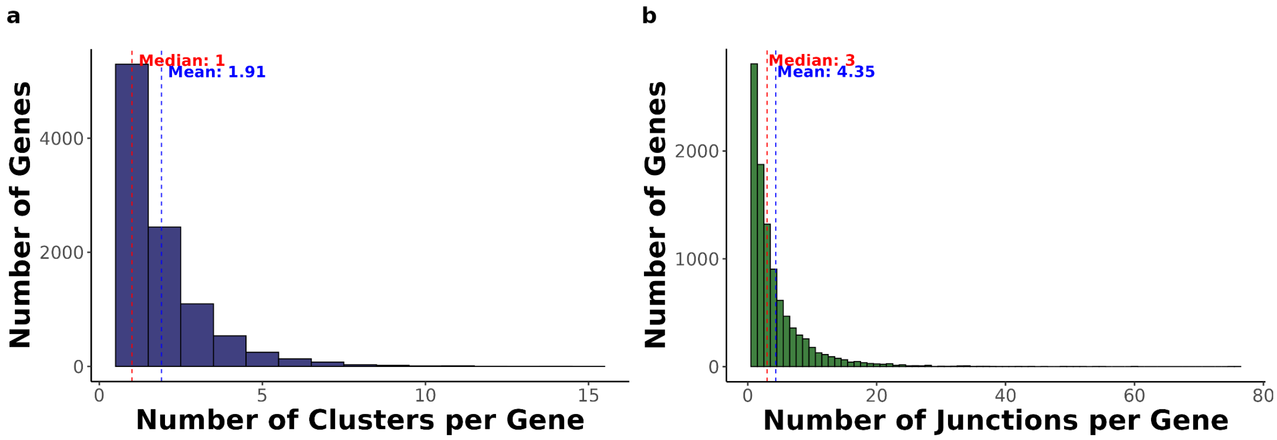


**Supplementary Fig. 1: Distribution of splice clusters and splice junctions per gene. (a)** Histogram showing the number of splice clusters assigned to each gene. The red dashed line indicates the median number of clusters per gene, and the blue dashed line indicates the mean. **(b)** Histogram showing the number of detected splice junctions assigned to each gene. The red and blue dashed lines indicate the median and mean number of junctions per gene, respectively.

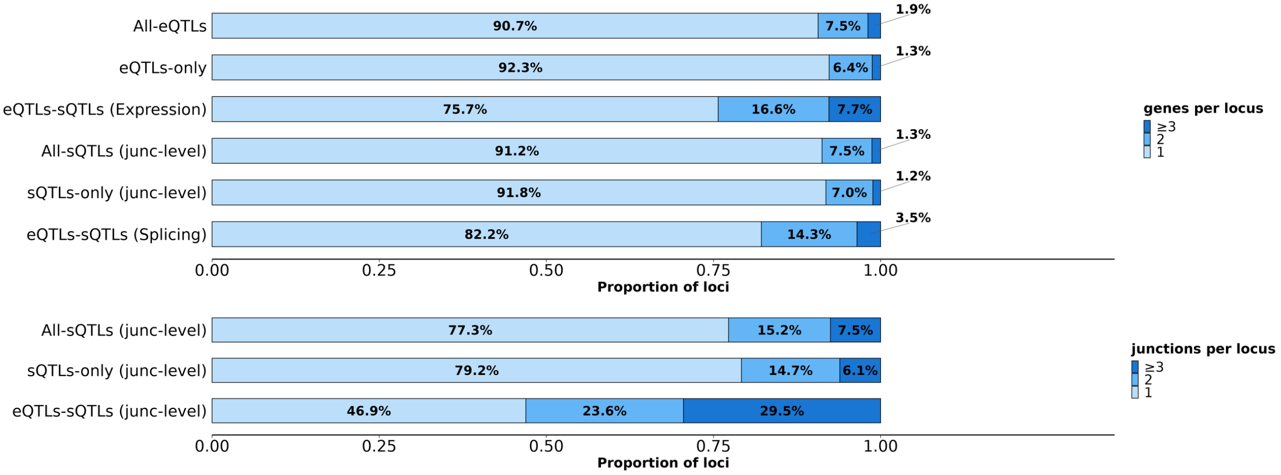


**Supplementary Fig. 2: Complexity of QTL loci at junction level.** Distribution of the number of genes or junctions per QTL locus. Stacked bars show the proportion of loci with one, two, or at least three target genes or junctions across all eQTLs, eQTL-only loci, overlapping eQTL–sQTL loci, all junction-level sQTLs, sQTL-only loci, and overlapping junction-level eQTL–sQTL loci.

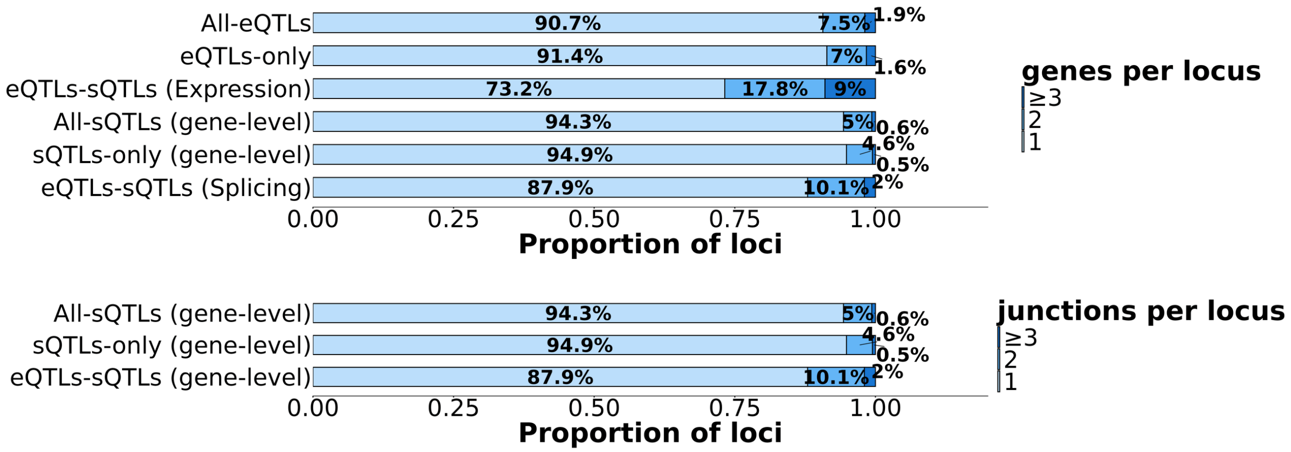


**Supplementary Fig. 3: Complexity of QTL loci at gene level.** Stacked bars show the proportion of QTL loci with one, two, or at least three target genes or junctions. The upper panel summarizes genes per locus across all eQTLs, eQTL-only loci, overlapping eQTL–sQTL expression signals, all gene-level sQTLs, sQTL-only loci, and overlapping eQTL–sQTL splicing signals. The lower panel summarizes junctions per locus for gene-level sQTL categories.

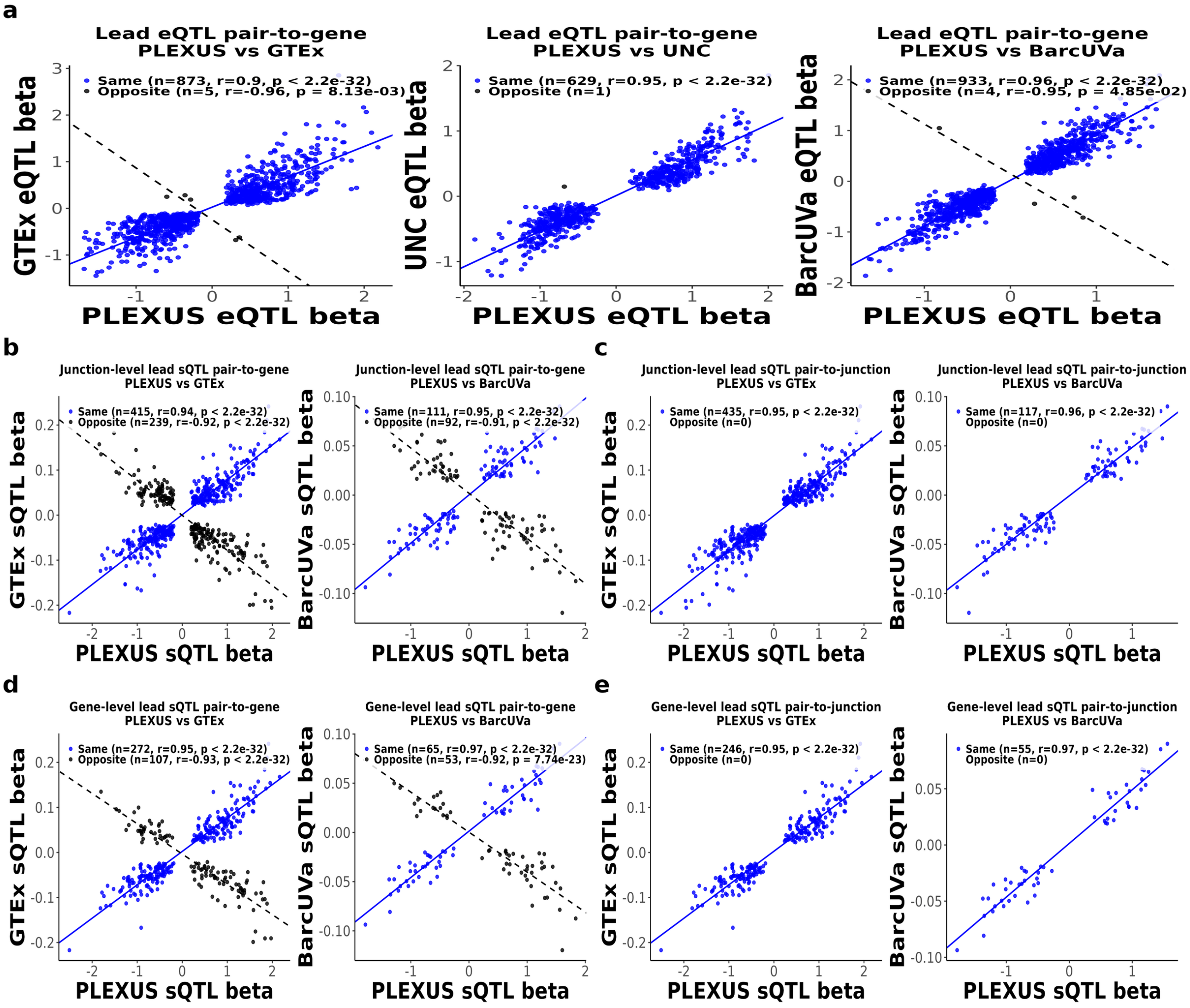


**Supplementary Fig. 4: Cross-cohort effect-size concordance of shared lead eQTL and sQTL associations.** **(a)** Effect size comparisons for exact lead-eQTL variant–gene pair matches between IBD Plexus and GTEx, UNC, or BarcUVa. **(b)** Effect size comparisons of junction-level lead sQTL– gene pair matches between IBD Plexus and GTEx or BarcUVa. **(c)** Effect size comparisons of junction-level lead sQTLs–junction pair matches after requiring an exact match of both the lead variant and splice junction. **(d)** Effect size comparisons of gene-level lead sQTLs after matching on lead variant and target gene. **(e)** Effect size comparisons of gene-level lead sQTLs after matching on lead variant and the associated splice junction. Effects were harmonized to the same effect allele before comparison. Blue points represent associations with concordant effect directions across cohorts, whereas black points represent associations with opposite effect directions. Solid blue and dashed black lines show separate linear fits for the concordant and opposite-direction subsets. Annotations report the number of matched associations, Pearson correlation coefficient $r$, and corresponding $P$ value.

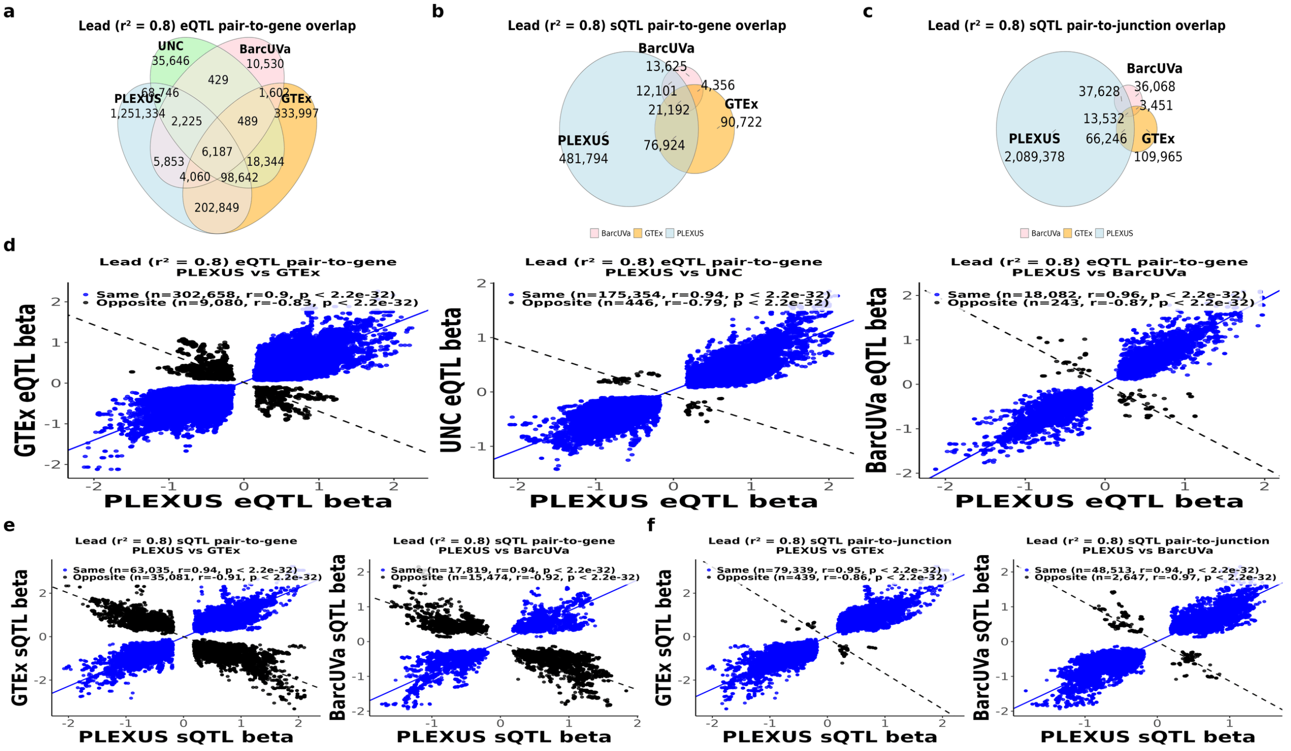


**Supplementary Fig. 5: Cross-cohort overlap and effect size concordance of LD-expanded QTL association sets.** For each cohort, lead-QTL association sets were expanded to include variants in strong linkage disequilibrium with the lead variant ($r^2 \geq 0.8$). **(a)** Euler diagram of lead-$r^2 \geq 0.8$ eQTL variant–gene pair overlap among IBD Plexus, GTEx, UNC, and BarcUVa. **(b)** Euler diagram of lead-$r^2 \geq 0.8$ sQTL variant–gene pair overlap among IBD Plexus, GTEx, and BarcUVa. **(c)** Euler diagram of lead-$r^2 \geq 0.8$ sQTL variant–junction pair overlap among the same three cohorts. Numbers in the Euler diagrams indicate the number of exact pair matches in each exclusive intersection. **(d)** Effect size concordance for shared eQTL variant–gene pairs between IBD Plexus and GTEx, UNC, or BarcUVa. **(e)** Effect size concordance for shared sQTL variant–gene pairs between IBD Plexus and GTEx or BarcUVa. **(f)** Effect size concordance for shared sQTL variant–junction pairs between IBD Plexus and GTEx or BarcUVa. Effects were oriented to the same allele before comparison. Blue points denote associations with concordant effect directions and black points denote associations with opposite effect directions. Solid blue and dashed black lines show separate linear fits for the concordant and opposite-direction subsets, respectively. Plot annotations report the number of associations, Pearson correlation coefficient $r$, and corresponding $P$ value for each subset.

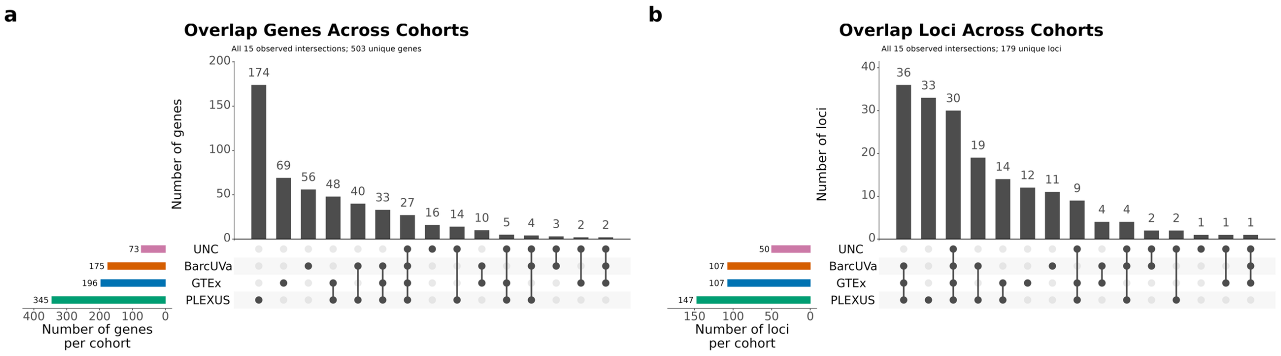


**Supplementary Fig. 6: Overlap of IBD GWAS-colocalized loci and QTL genes across colon QTL resources.** **(a)** UpSet plot showing overlap of IBD GWAS-colocalized loci across IBD Plexus eQTLs, IBD Plexus sQTLs, GTEx eQTLs, GTEx sQTLs, BarcUVa eQTLs, BarcUVa sQTLs, and UNC eQTLs. Horizontal bars show the size of each input set, vertical bars show intersection sizes, and connected black dots indicate the datasets contributing to each intersection. **(b)** UpSet plot showing overlap of IBD GWAS-colocalized QTL genes across the same datasets and QTL classes. This analysis summarizes both dataset-specific and shared colocalized genes across expression and splicing QTL resources.

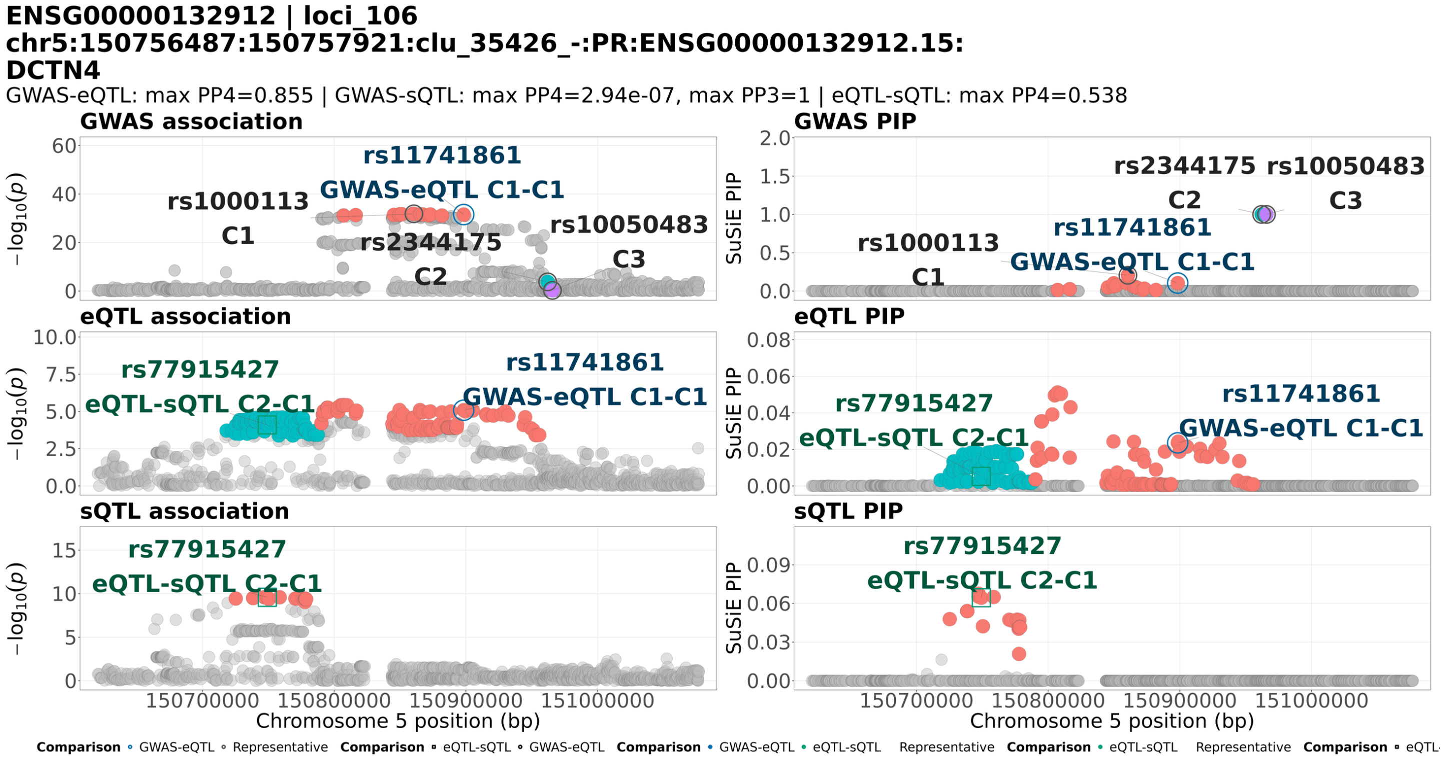


**Supplementary Fig. 7: Distinct eQTL components colocalize with GWAS and splicing at *DCTN4*.** Regional association statistics ($-\log_{10} P$, left) and SuSiE posterior inclusion probabilities (PIPs, right) are shown for IBD GWAS, *DCTN4* expression, and usage of junction `chr5:150756487:150757920` in LeafCutter2 cluster `clu_35426_-` (top to bottom). Each point represents a variant. Credible-set variants are colored by component within each trait, with C1, C2, and C3 shown in salmon, turquoise, and purple, respectively. Gray points indicate other variants. Blue circles highlight `rs11741861`, a candidate shared variant for GWAS component 1 and eQTL component 1. Green squares highlight `rs77915427`, a candidate shared variant for eQTL component 2 and sQTL component 1. Additional labeled variants mark representative variants of individual components. Labels such as C2–C1 indicate the component pair in the order of the named traits. Maximum PP4 values were 0.855 for GWAS–eQTL and 0.538 for eQTL–sQTL. GWAS–sQTL comparisons supported distinct associations (maximum PP3 = 1.000, maximum PP4 = $2.94 \times 10^{-7}$). This example illustrates how separate expression components can share associations with IBD and splicing while the GWAS and sQTL signals remain distinct.

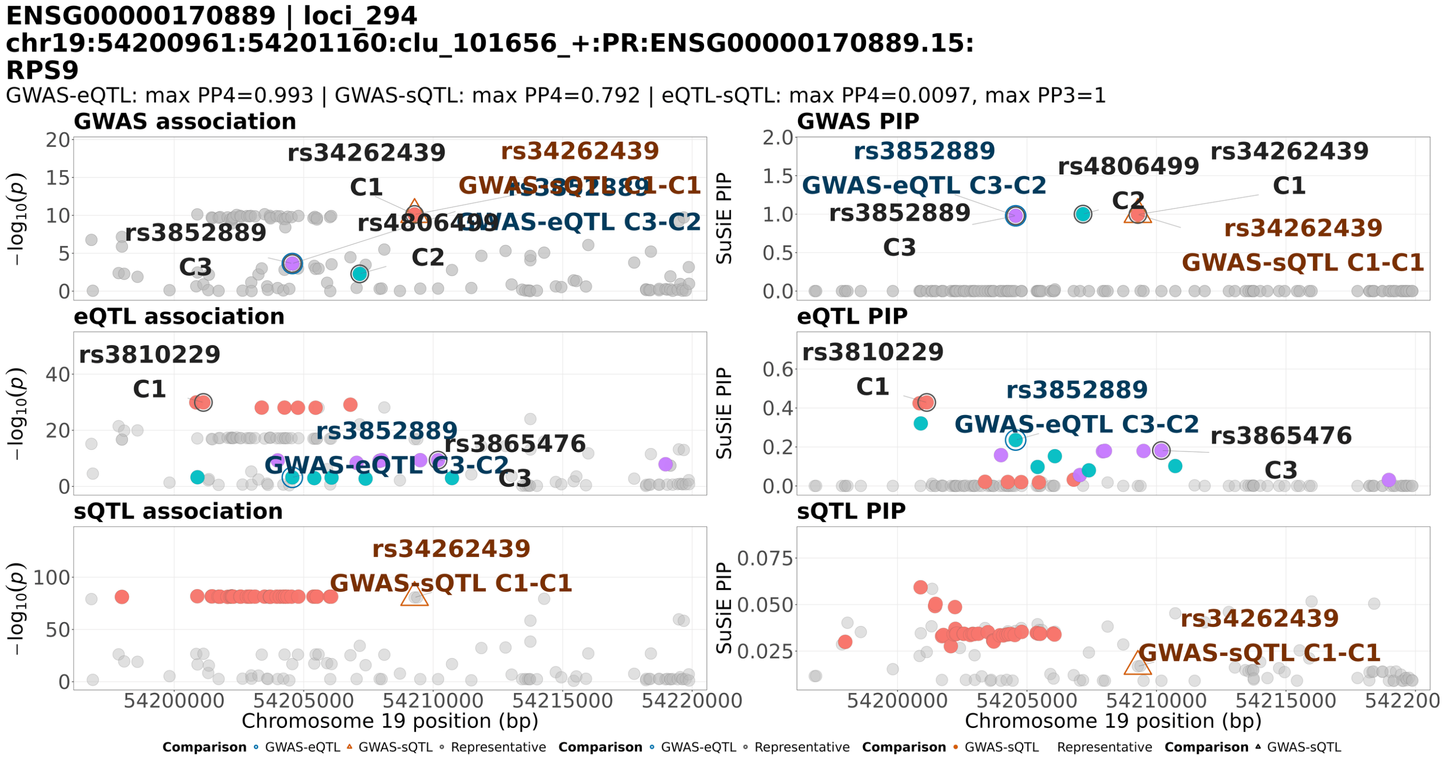


**Supplementary Fig. 8: Distinct GWAS components colocalize with expression and splicing at *RPS9*.** Regional association statistics ($-\log_{10} P$, left) and SuSiE posterior inclusion probabilities (PIPs, right) are shown for IBD GWAS, *RPS9* expression, and usage of junction `chr19:54200961:54201159` in LeafCutter2 cluster `clu_101656_+` (top to bottom). Each point represents a variant. Credible-set variants are colored by component within each trait, with C1, C2, and C3 shown in salmon, turquoise, and purple, respectively. Gray points indicate other variants. Blue circles highlight `rs3852889`, a candidate shared variant for GWAS component 3 and eQTL component 2. Orange triangles highlight `rs34262439`, a candidate shared variant for GWAS component 1 and sQTL component 1. Additional labeled variants mark representative variants of individual components. Labels such as C3–C2 indicate the component pair in the order of the named traits. Maximum PP4 values were 0.993 for GWAS–eQTL and 0.792 for GWAS–sQTL, whereas eQTL–sQTL comparisons showed little support for colocalization (maximum PP4 = 0.0097) and strong support for distinct associations (maximum PP3 = 1.000). This example illustrates how expression and splicing can colocalize with different GWAS components without supporting colocalization with each other.

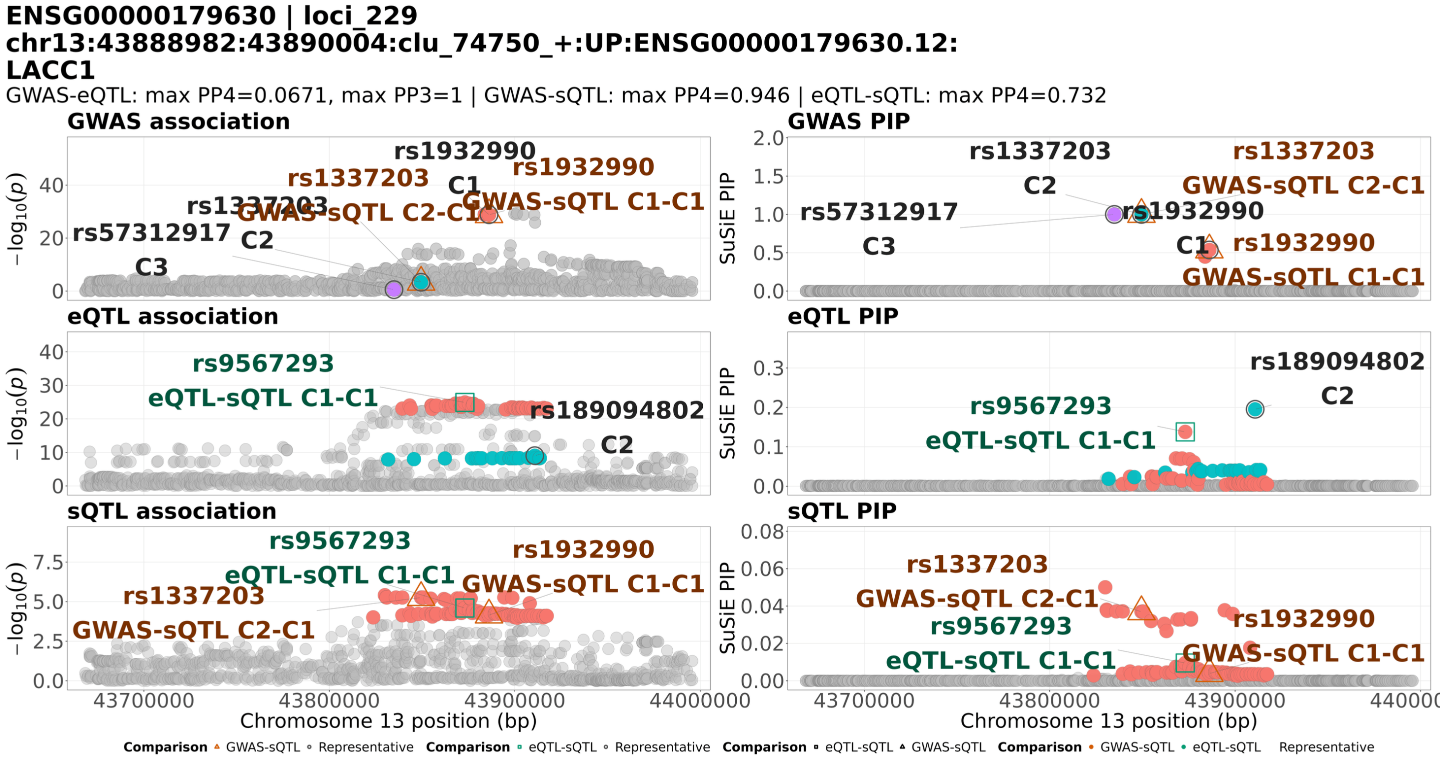


**Supplementary Fig. 9: Pairwise colocalization and limited sQTL fine-mapping resolution at *LACC1*.** Regional association statistics ($-\log_{10} P$, left) and SuSiE posterior inclusion probabilities (PIPs, right) are shown for IBD GWAS, *LACC1* expression, and usage of junction `chr13:43888982:43890003` in LeafCutter2 cluster `clu_74750_+` (top to bottom). Each point represents a variant. Credible-set variants are colored by component within each trait, with C1, C2, and C3 shown in salmon, turquoise, and purple, respectively. Gray points indicate other variants. Orange triangles highlight candidate shared variants for GWAS–sQTL comparisons, including `rs1932990` for GWAS component 1 and sQTL component 1, and `rs1337203` for GWAS component 2 and sQTL component 1. Green squares highlight `rs9567293`, a candidate shared variant for eQTL component 1 and sQTL component 1. Additional labeled variants mark representative variants of individual components. Component-pair labels follow the order of the named traits. Maximum PP4 values were 0.946 for GWAS–sQTL and 0.732 for eQTL–sQTL, whereas GWAS–eQTL comparisons favored distinct associations (maximum PP3 = 1.000, maximum PP4 = 0.0671). The sQTL component assigns low inclusion probabilities across many variants, illustrating limited fine-mapping resolution when the same sQTL component supports colocalization with different GWAS and eQTL components through different candidate shared variants.

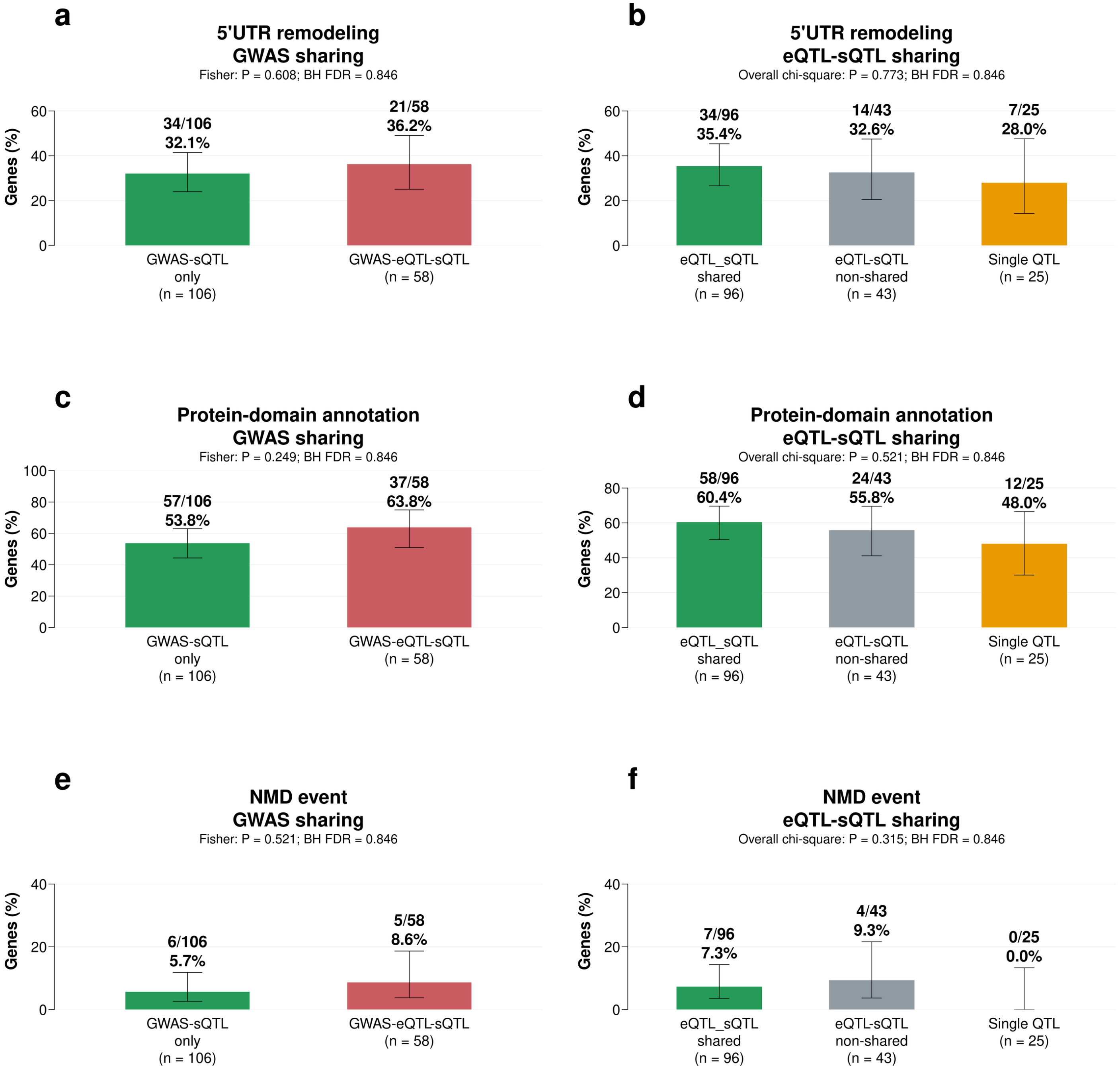


**Supplementary Fig. 10: Candidate splicing mechanisms among IBD GWAS-colocalized sGenes stratified by GWAS and molecular QTL sharing.** Proportions of genes with candidate 5′ UTR remodeling events **(a,b)**, protein-domain-associated splicing **(c,d)**, or nonsense-mediated decay (NMD) events **(e,f)**. **(a,c,e)** Genes with only sQTL–GWAS colocalization ($n = 106$) were compared with genes with both eQTL–GWAS and sQTL–GWAS colocalization ($n = 58$). **(b,d,f)** Genes were grouped according to whether their eQTL and sQTL signals colocalized (shared; $n = 96$), did not meet the colocalization threshold (non-shared; $n = 43$), or only an sQTL was detected (single QTL; $n = 25$). Bars show the percentage of genes with each candidate mechanism; labels indicate the corresponding gene counts, group totals, and percentages. Differences between groups were assessed using Fisher's exact test for GWAS-sharing comparisons and an overall chi-square test for eQTL–sQTL-sharing comparisons. Raw $P$ values and Benjamini–Hochberg (BH)-adjusted $P$ values are displayed above each panel. These annotations represent candidate mechanisms rather than experimentally validated functional effects.

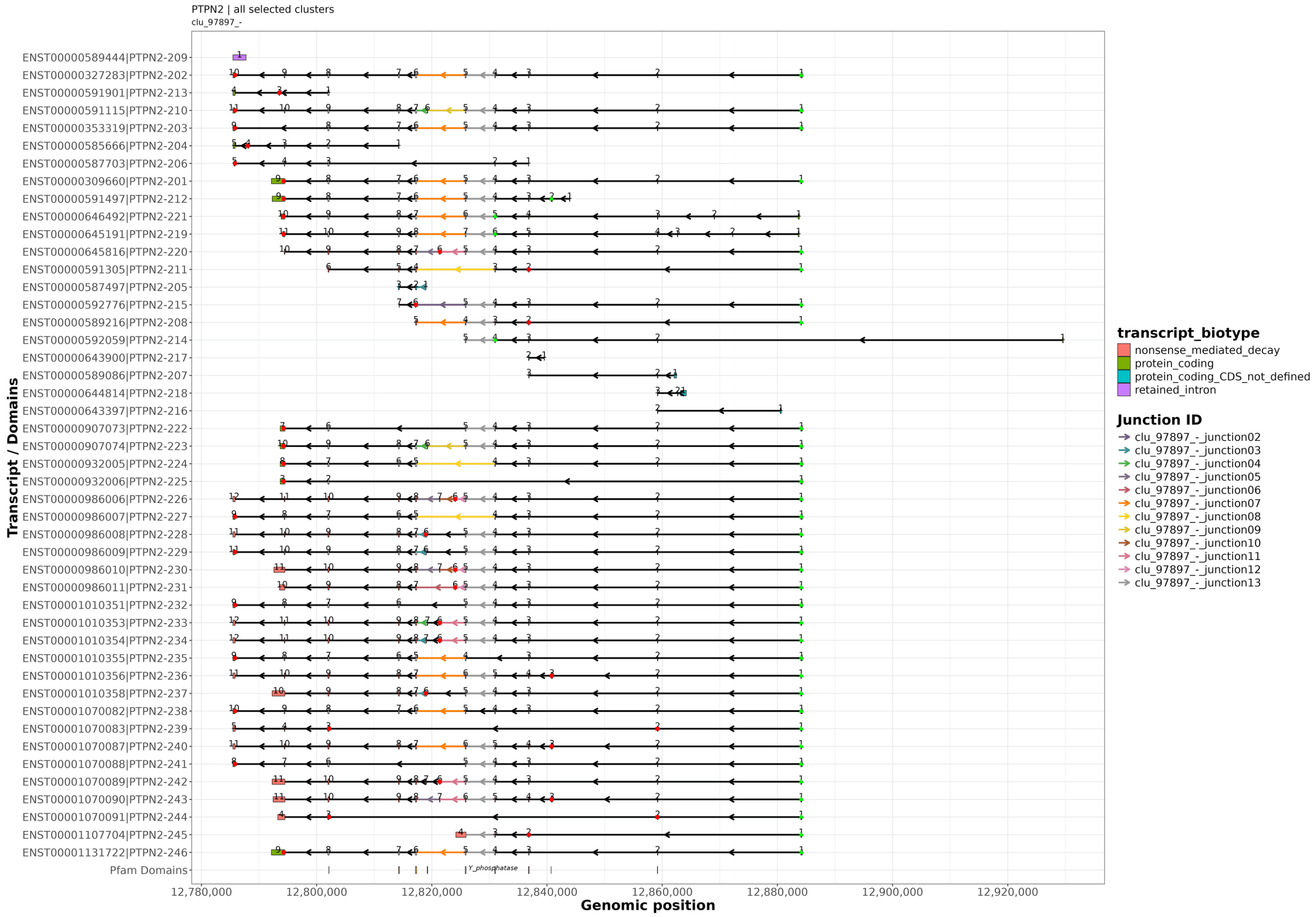


**Supplementary Fig. 11: GENCODE transcript models and alternative splice junctions of *PTPN2*.** Overview of GENCODE-annotated *PTPN2* transcripts showing alternative splice junctions in the LeafCutter2-defined cluster `clu_97897_-`. Transcript identifiers and names are displayed on the left. Boxes represent exons, with exon numbers indicated within each transcript, and connecting lines represent introns. Exons are colored by transcript annotation as protein coding, nonsense-mediated decay (NMD), protein coding with an undefined coding sequence (CDS), or retained intron. Colored connections identify cluster junctions represented in the annotated transcript models, with junction identifiers listed on the right. Arrows indicate the negative-strand direction of transcription. Genomic positions of the Pfam protein tyrosine phosphatase domain, Y_phosphatase, are shown below the transcript models. The x-axis shows chromosome 18 coordinates.

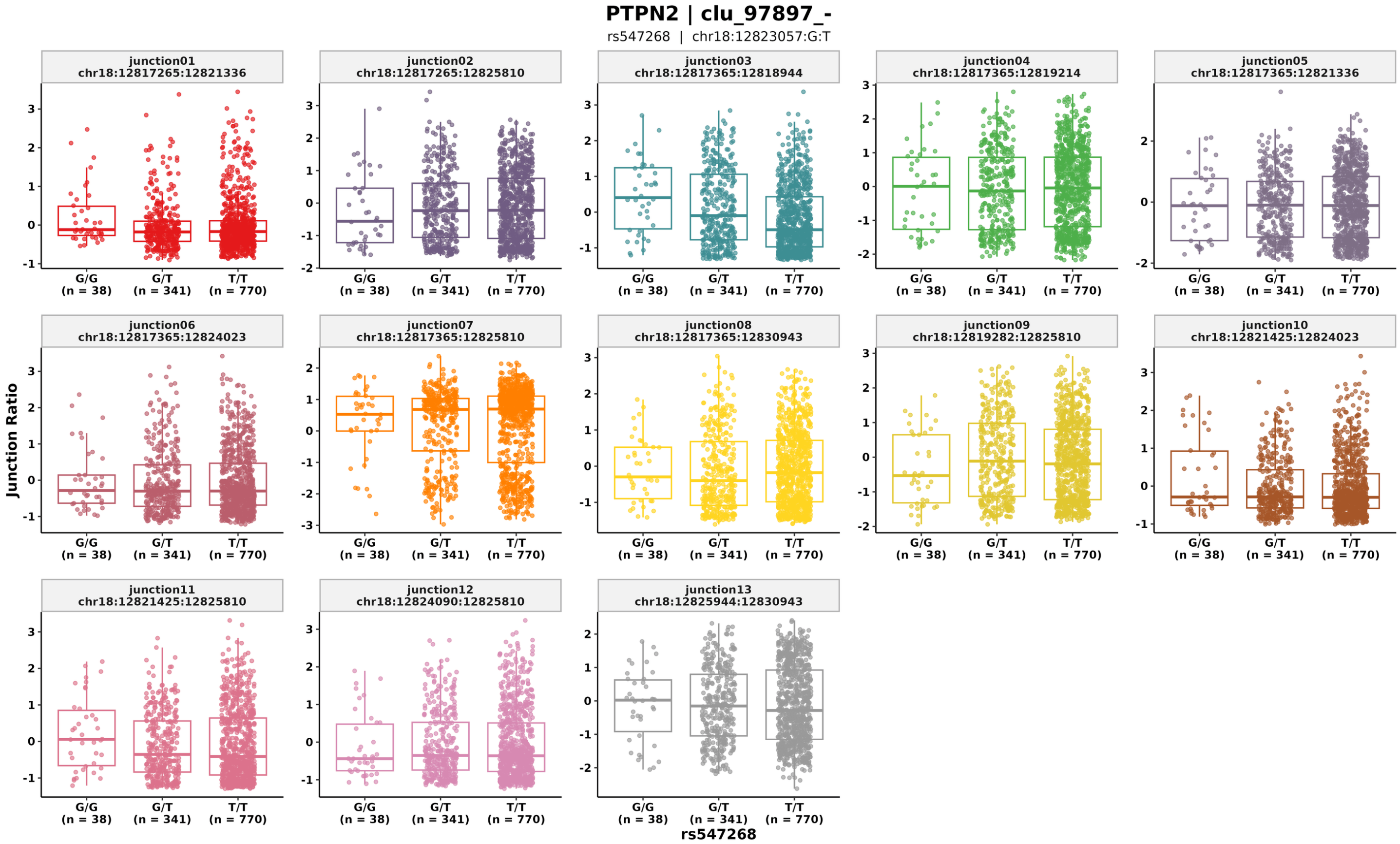


**Supplementary Fig. 12: Genotype-associated splice junction usage in *PTPN2*.** Normalized usage of 13 junctions in the LeafCutter2-defined cluster `clu_97897_-`, stratified by GWAS index variant `rs547268` genotype. Junction identifiers and chromosome 18 coordinates are displayed above each plot. Colors distinguish individual junctions and correspond to those used for the junctions represented in the transcript models. Individual samples are superimposed on the box plots, and sample counts are shown below each genotype: G/G, $n = 38$; G/T, $n = 341$; and T/T, $n = 770$.

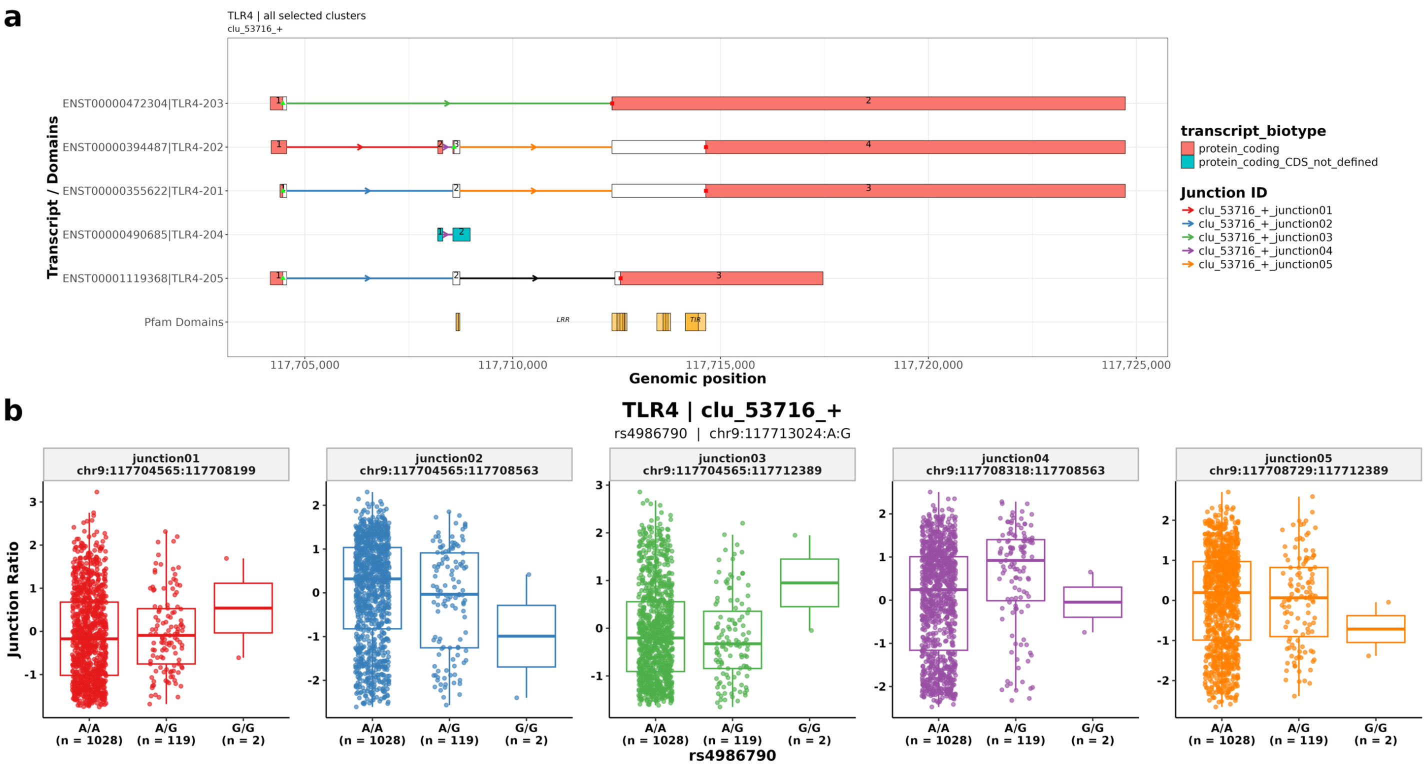


**Supplementary Fig. 13: Transcript models and genotype-associated splice junction usage of *TLR4*.** **(a)** GENCODE transcript models showing alternative splice junctions in the LeafCutter2-defined cluster `clu_53716_+`. Exons are numbered within each transcript and colored by transcript annotation as protein coding or protein coding with an undefined coding sequence (CDS). Colored connections identify cluster junctions, and arrows indicate the direction of transcription. Genomic positions of the Pfam leucine-rich repeat (LRR) and Toll/interleukin-1 receptor (TIR) domains are shown below the transcript models. **(b)** Normalized usage of the five junctions in this cluster stratified by `rs4986790` genotype. Junction identifiers and chromosome 9 coordinates are displayed above each plot, with junction colors corresponding to those in panel (a). Individual samples are superimposed on the box plots, and sample counts are shown below each genotype: A/A, $n$ = 1,028; A/G, $n$ = 119; and G/G, $n$ = 2.

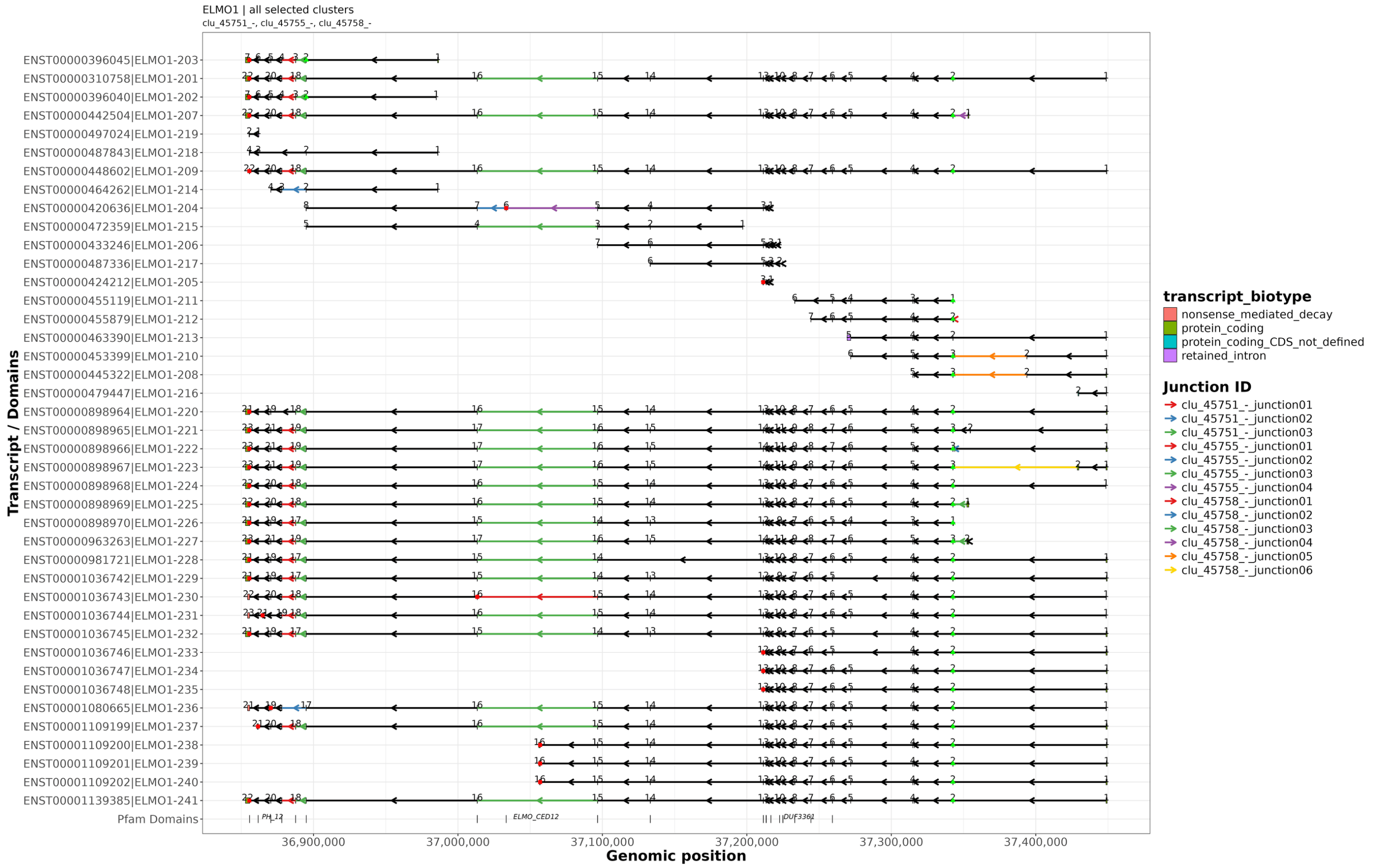


**Supplementary Fig. 14: GENCODE transcript models of *ELMO1*.** Overview of GENCODE-annotated *ELMO1* transcripts across the genomic region containing the LeafCutter2-defined clusters `clu_45751_-`, `clu_45755_-`, and `clu_45758_-`. Transcript identifiers and names are displayed on the left. Boxes represent exons, with exon numbers indicated within each transcript, and connecting lines represent introns. Exons are colored by transcript annotation as protein coding, nonsense-mediated decay (NMD), protein coding with an undefined coding sequence (CDS), or retained intron. Arrows indicate the negative-strand direction of transcription. Genomic positions of the Pfam domains PH_12, ELMO_CED12, and DUF3361 are shown below the transcript models. The x-axis shows chromosome 7 coordinates.

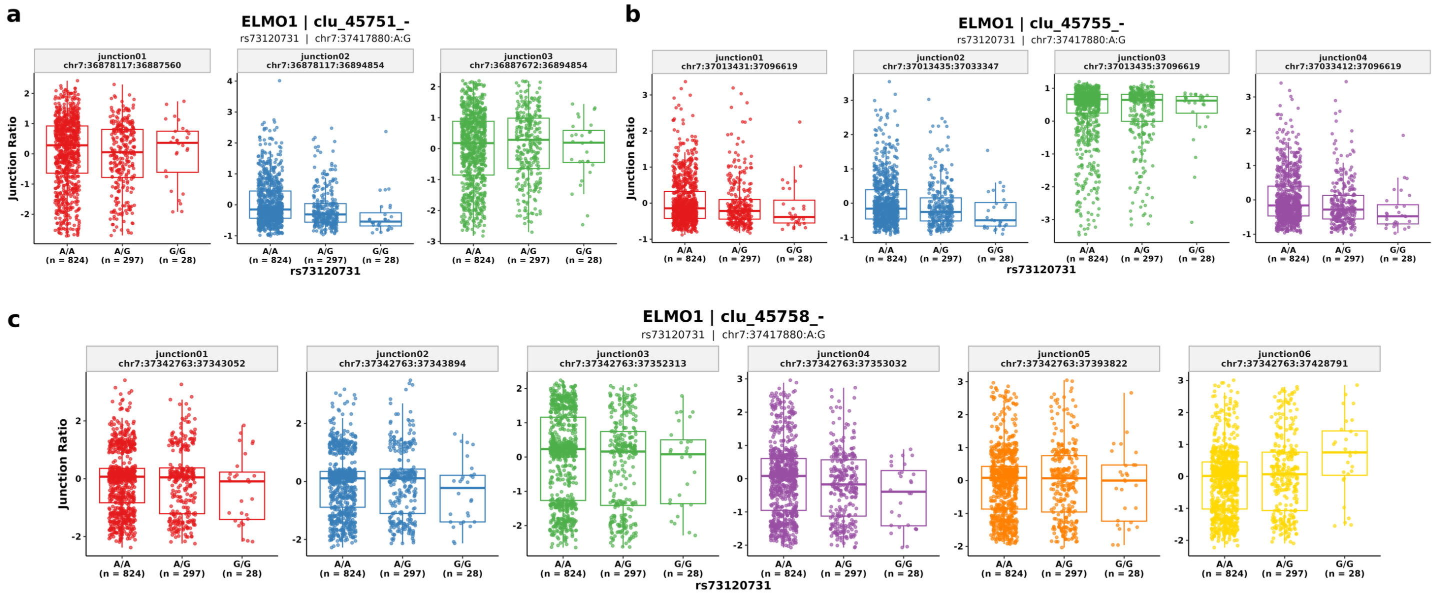


**Supplementary Fig. 15: Genotype-associated splice junction usage across three *ELMO1* splice clusters.** Normalized junction usage in the LeafCutter2-defined clusters **(a)** `clu_45751_-`, **(b)** `clu_45755_-`, and **(c)** `clu_45758_-`, stratified by `rs73120731` genotype. Panels show three, four, and six junctions, respectively. Junction identifiers and chromosome 7 coordinates are displayed above each plot. Colors distinguish junctions within each cluster. Individual samples are superimposed on the box plots, and sample counts are shown below each genotype: A/A, $n = 824$; A/G, $n = 297$; and G/G, $n = 28$.

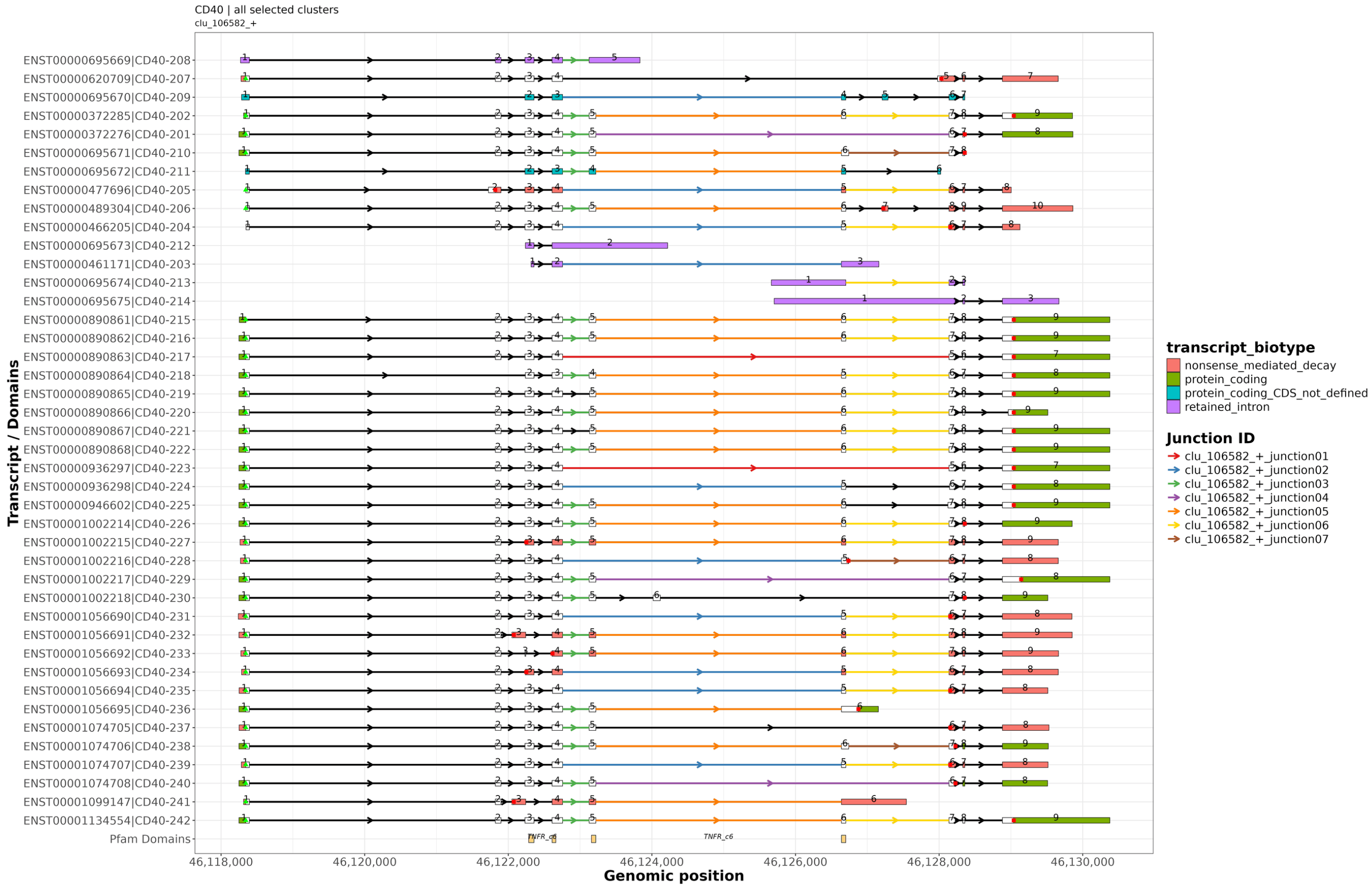


**Supplementary Fig. 16: GENCODE transcript models and alternative splice junctions of *CD40*.** Overview of GENCODE-annotated *CD40* transcripts showing alternative splice junctions in the LeafCutter2-defined cluster `clu_106582_+`. Transcript identifiers and names are displayed on the left. Boxes represent exons, with exon numbers indicated within each transcript, and connecting lines represent introns. Exons are colored by transcript annotation as protein coding, nonsense-mediated decay (NMD), protein coding with an undefined coding sequence (CDS), or retained intron. Colored connections identify cluster junctions, with junction identifiers listed on the right. Arrows indicate the positive-strand direction of transcription. Genomic positions of the Pfam TNFR_c6 domain are shown below the transcript models. The x-axis shows chromosome 20 coordinates.

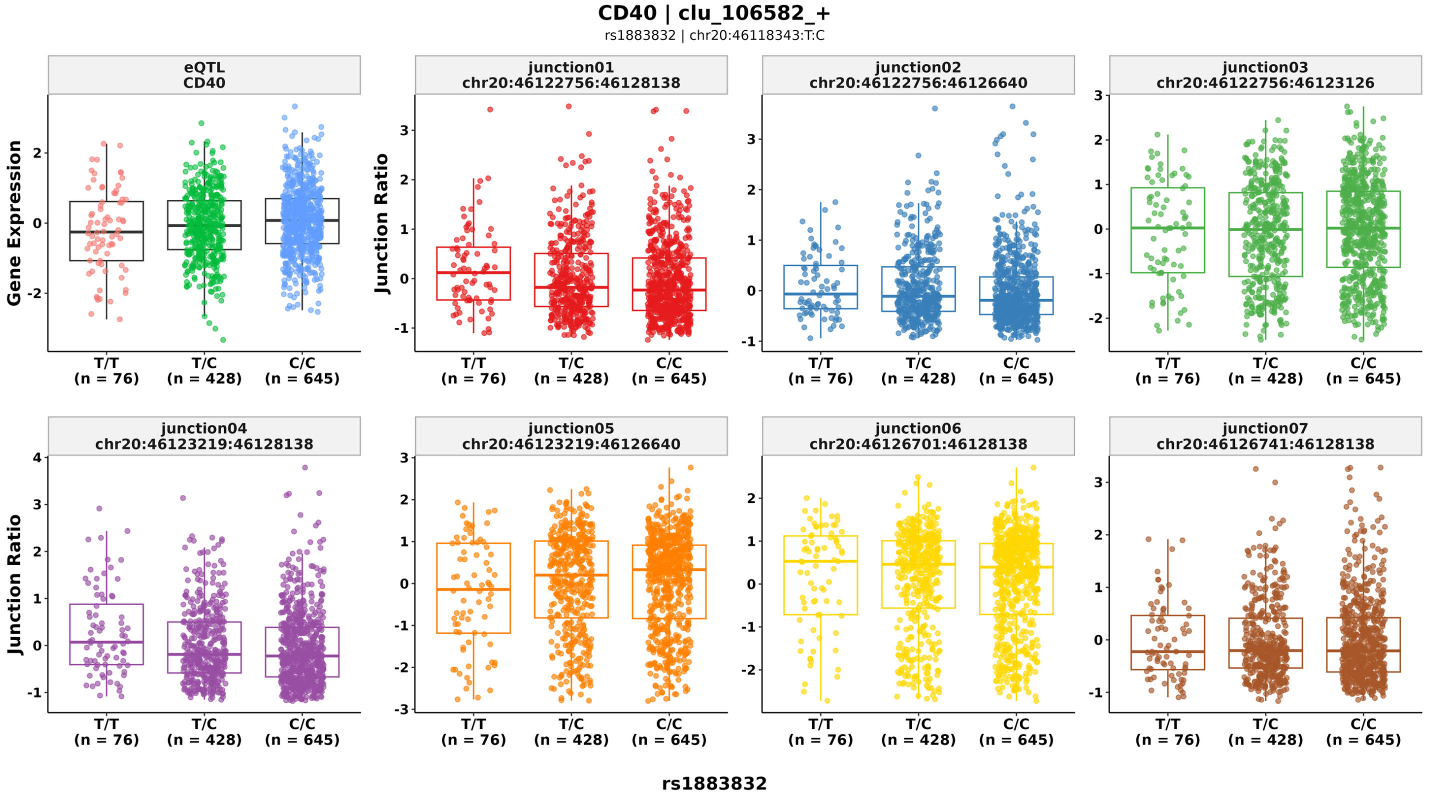


**Supplementary Fig. 17: Genotype-associated gene expression and splice junction usage in *CD40*.** Normalized *CD40* expression and usage of seven junctions in the LeafCutter2 defined cluster `clu_106582_+`, stratified by `rs1883832` genotype. The upper-left panel shows gene expression, with points colored by genotype. The remaining panels show individual splice junctions, with junction identifiers and chromosome 20 coordinates displayed above each plot. Junction colors correspond to those used in the transcript models. Individual samples are superimposed on the box plots, and sample counts are shown below each genotype: T/T, $n = 76$; T/C, $n = 428$; and C/C, $n = 645$. Boxes indicate the interquartile range, horizontal lines indicate medians, and whiskers extend to the most extreme observations within 1.5 times the interquartile range of the box boundaries.

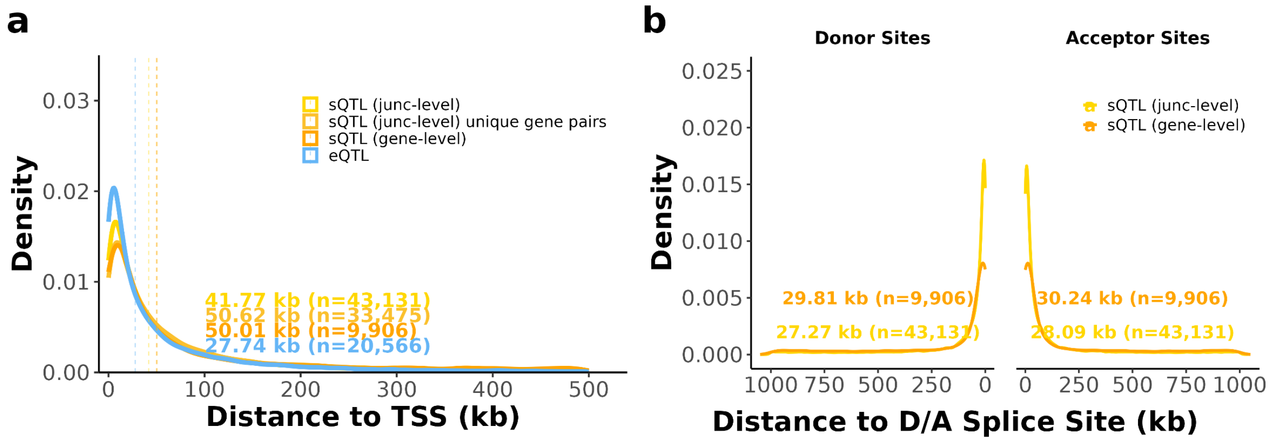


**Supplementary Fig. 18: Genomic distances of lead eQTL and sQTL variants to transcription start sites and splice sites. (a)** Density distributions of absolute distances between lead QTL variants and the transcription start site (TSS) of their associated genes, comparing eQTLs, gene-level sQTLs, junction-level sQTLs, and junction-level sQTLs collapsed to unique variant–gene pairs. Dashed vertical lines indicate median distances, with corresponding values and sample sizes shown. **(b)** Density distributions of distances between lead sQTL variants and splice donor and acceptor sites, comparing gene-level and junction-level sQTLs. Median distances and sample sizes are indicated for each group.

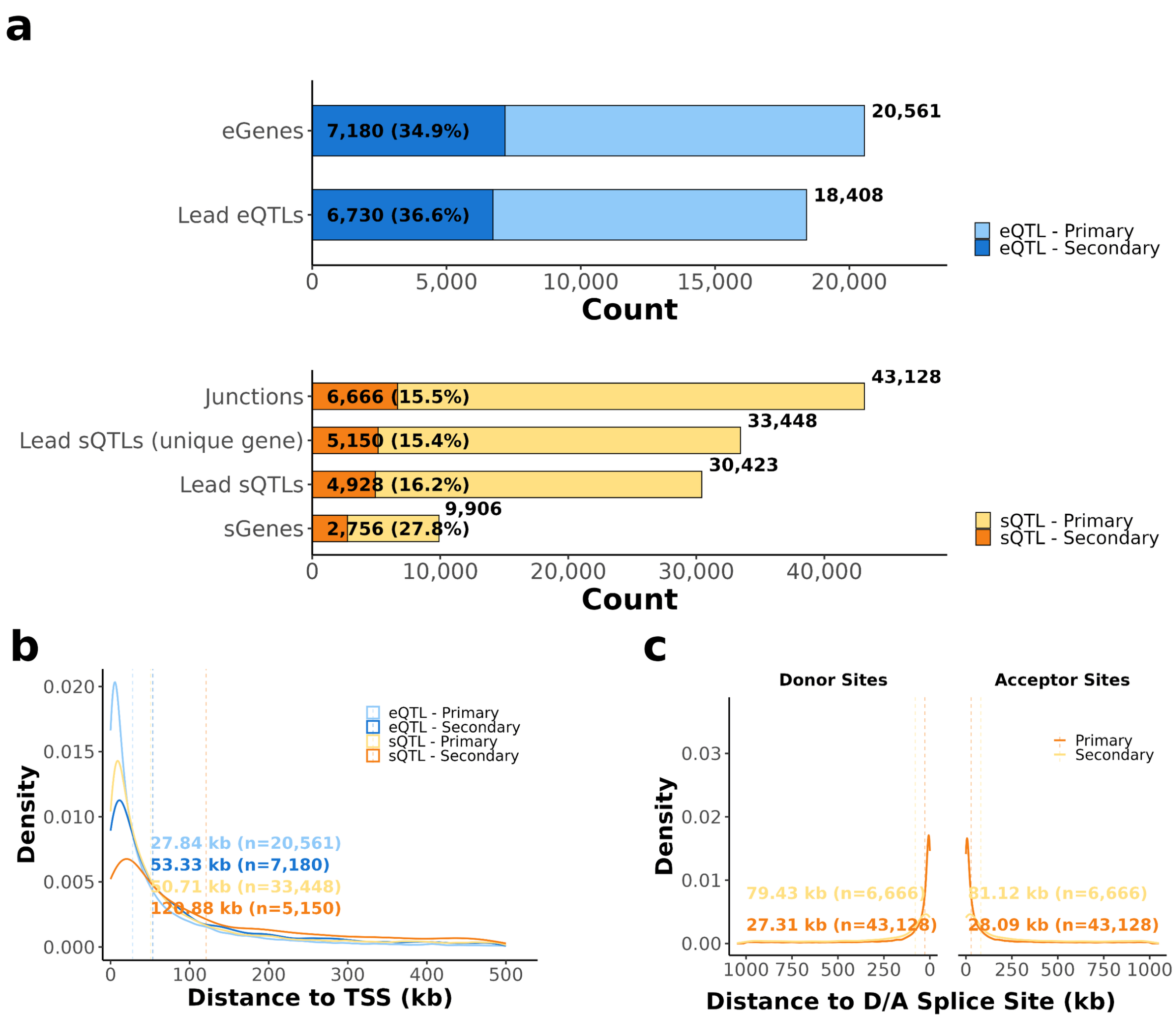


**Supplementary Fig. 19: Primary and secondary QTL signals differ in abundance and genomic proximity.** **(a)** Stacked horizontal bars summarize the number of primary and secondary eQTL and sQTL signals after conditional analysis. Expression signals are shown for eGenes and lead eQTLs, whereas splicing signals are shown for junctions, lead sQTLs collapsed to unique genes, lead sQTLs, and sGenes. Bar labels indicate the number and percentage of secondary signals, and totals are shown at the right of each bar. **(b)** Density distributions of distances from primary and secondary eQTL and sQTL variants to the nearest transcription start site (TSS). Dashed vertical lines indicate group-specific median distances. **(c)** Density distributions of distances from primary and secondary sQTL variants to the nearest splice donor or acceptor site. Both primary and secondary sQTL signals are concentrated near canonical splice sites, with dashed vertical lines indicating median distances.

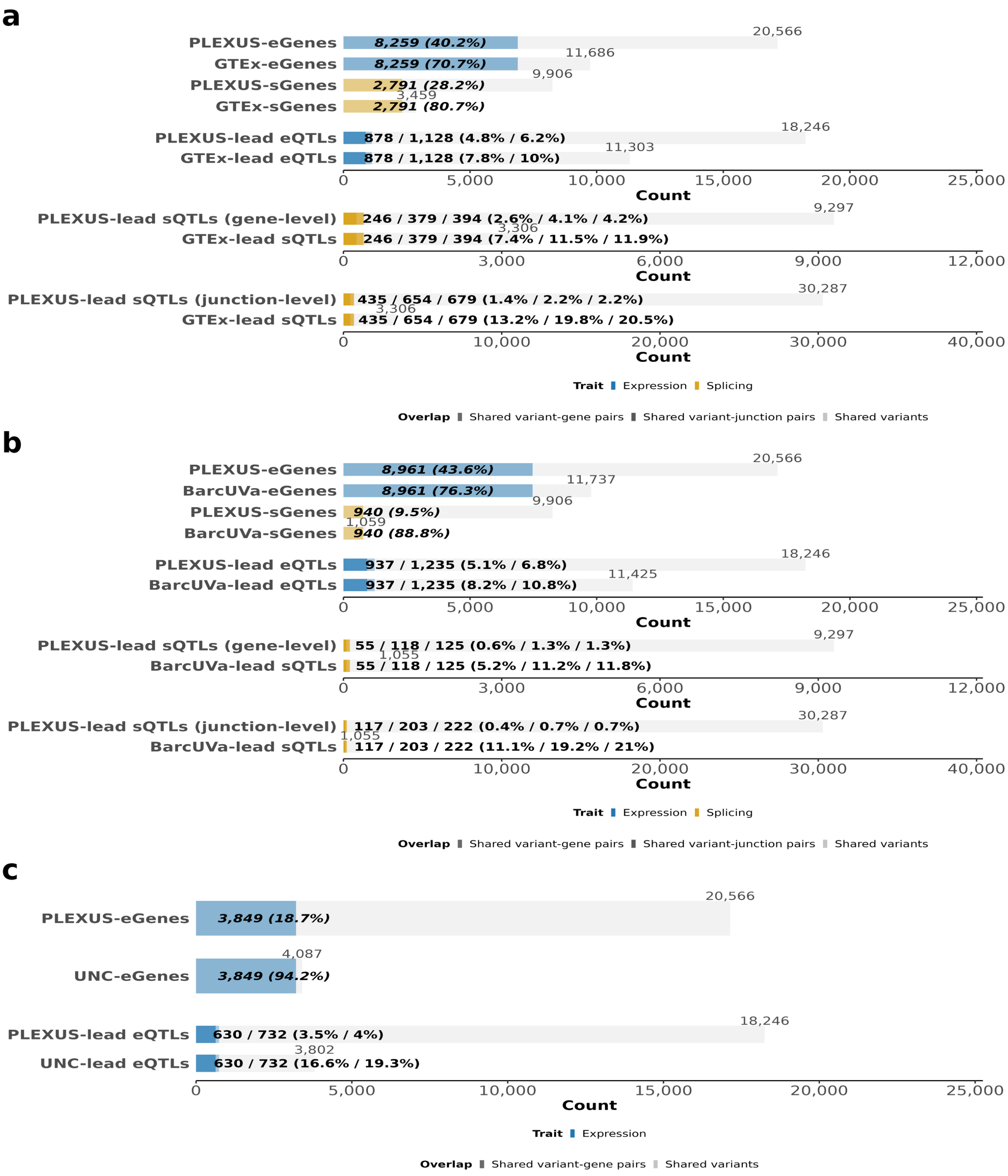


**Supplementary Fig. 20: Pairwise overlap of QTL genes and lead QTL associations between IBD Plexus and external cohorts.** **(a)** Overlap between IBD Plexus and GTEx. **(b)** Overlap between IBD Plexus and BarcUVa. **(c)** Overlap between IBD Plexus and UNC. Gray bars show the total number of genes or distinct lead variants in each cohort, and colored portions indicate cross-cohort overlap. For eGene and sGene rows, the label gives the number and percentage of genes detected in both cohorts. For lead-eQTL rows, the two values separated by a slash represent the number of shared variant–gene pairs and the number of shared lead variants, respectively. For gene-level and junction-level lead-sQTL rows, the three values represent shared variant–gene pairs, shared variant–junction pairs, and shared lead variants, respectively. Percentages are reported in the same order and are calculated relative to the total shown for the corresponding cohort row. The overlap numerator is identical for the two cohort rows, whereas the percentage differs according to the cohort-specific denominator. Gene-level and junction-level lead-sQTL sets are displayed separately. The UNC comparison is restricted to expression QTLs.

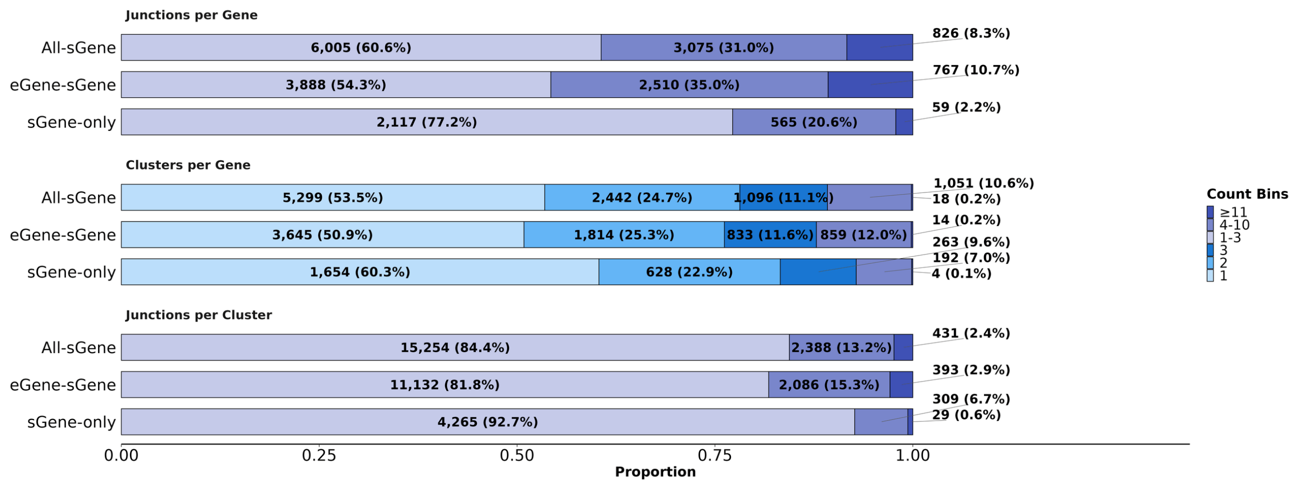


**Supplementary Fig. 21: Genomic complexity across sGene–only genes, eGene–only genes, and shared eGene-sGene genes..** Complexity of splicing targets among all sGenes, eGene–sGene genes, and sGene-only genes. Stacked bars show the distributions of junctions per gene, clusters per gene, and junctions per cluster, with segments representing count bins.

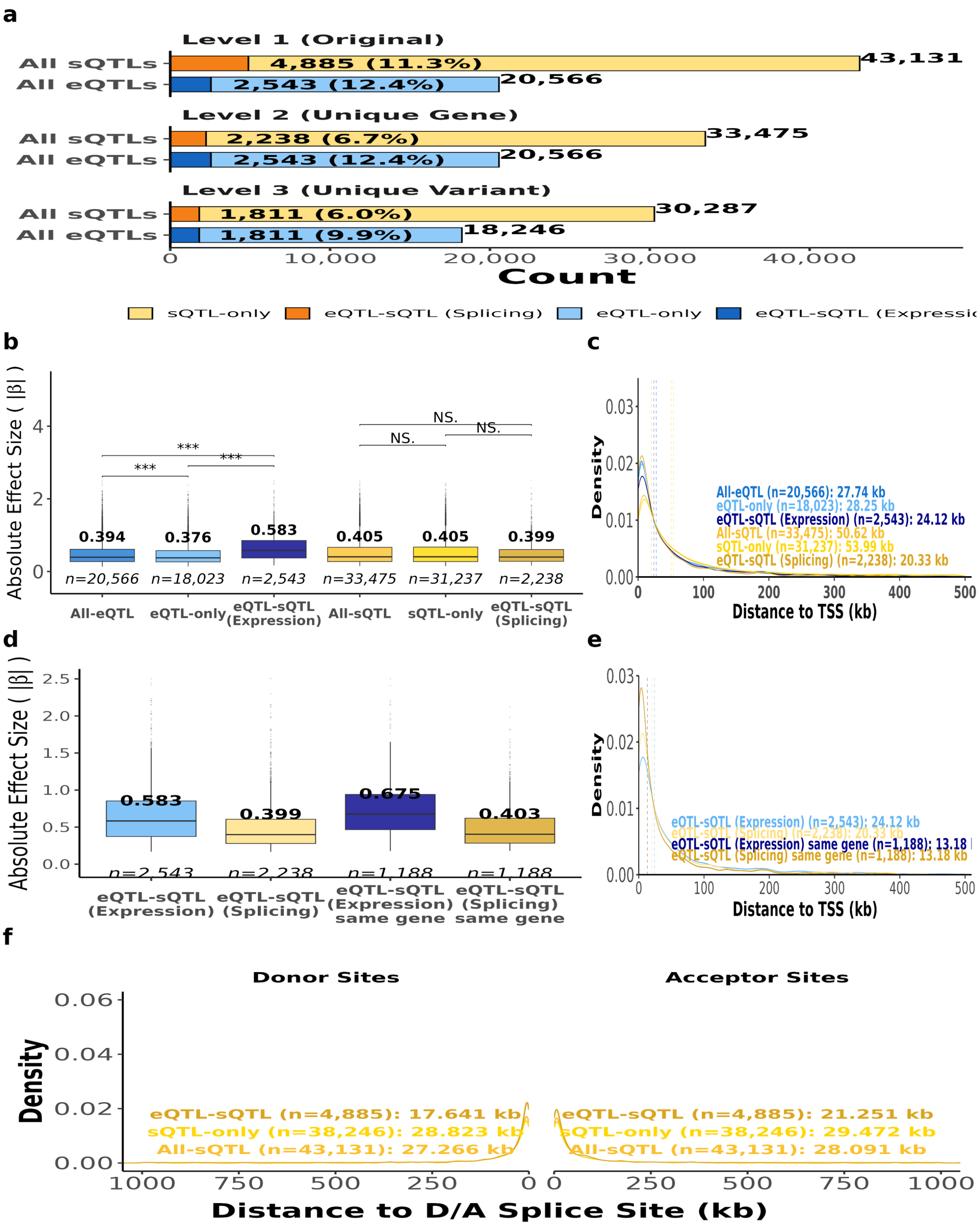


**Supplementary Fig. 22: Shared lead eQTLs and junction-level sQTLs: overlap, effect sizes, and distances to transcription start and splice sites.** **(a)** Trait-specific and shared lead-QTL counts at three resolutions. Level 1 retains all original associations, including distinct junctions and target genes. Level 2 retains unique lead variant–gene pairs, selecting the junction with the largest absolute splicing effect when multiple junctions map to the same pair. Level 3 retains distinct variants, selecting the variant–gene association with the largest absolute effect. Expression-side and splicing-side overlap counts may differ at levels 1 and 2 but are identical at level 3, where each shared variant is counted once. **(b)** Absolute effect sizes ($|\beta|$) for all, trait-specific, and shared associations, shown separately for expression and splicing at level 2. Bold values indicate medians; italicized values indicate association counts. **(c)** Absolute distances from lead variants to the associated genes' transcription start sites (TSSs) for the groups in **(b)**. Dotted lines and annotations indicate medians and sample sizes. **(d)** Absolute expression and splicing effect sizes for shared associations at level 2 before and after requiring both QTL types to target the same gene. **(e)** TSS distances for the shared associations in **(d)**. **(f)** Distances from junction-level lead sQTLs to associated donor and acceptor sites for all, sQTL-only, and shared associations at level 1. Donor distances are plotted left of zero and acceptor distances right; annotations give median absolute distances and sample sizes. Throughout, "eQTL–sQTL (Expression)" and "eQTL–sQTL (Splicing)" denote the expression-side and splicing-side associations of variants represented in both traits. Brackets indicate pairwise tests; NS, not significant; $^{**}P < 0.01$; $^{***}P < 0.001$.

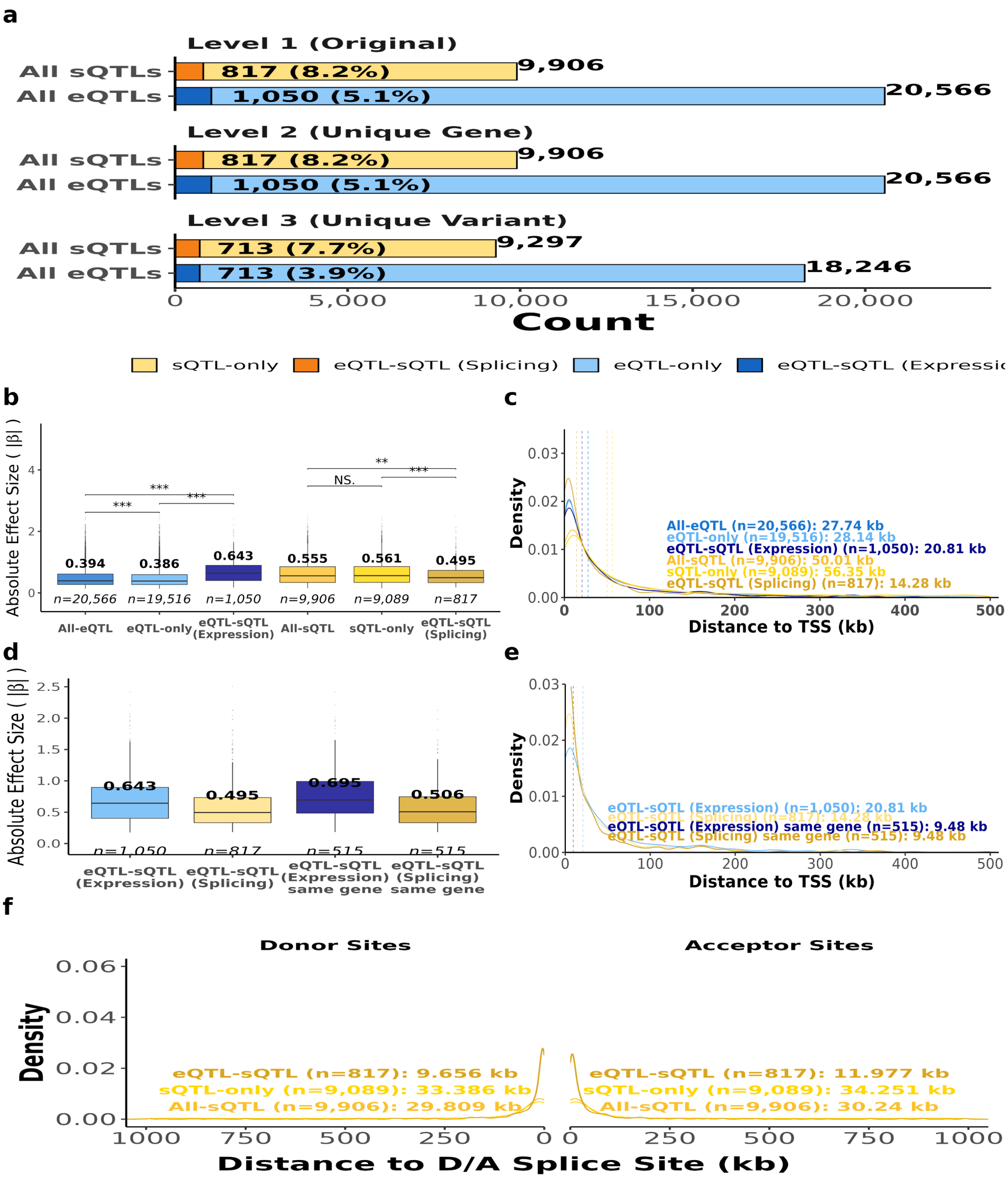


**Supplementary Fig. 23: Shared lead eQTLs and gene-level sQTLs: overlap, effect sizes, and distances to transcription start and splice sites.** **(a)** Trait-specific and shared lead-QTL counts at three resolutions. Level 1 retains original gene-level associations, including variants associated with multiple genes. Level 2 retains unique lead variant–gene pairs. Levels 1 and 2 are identical for gene-level sQTLs because each original record is already defined at the gene level. Level 3 retains distinct variants, selecting the variant–gene association with the largest absolute effect. Expression-side and splicing-side overlap counts may differ before variant-level deduplication but are identical at level 3, where each shared variant is counted once. **(b)** Absolute effect sizes ($|\beta|$) for all, trait-specific, and shared associations, shown separately for expression and splicing at level 2. Bold values indicate medians; italicized values indicate association counts. **(c)** Absolute distances from lead variants to the associated genes' transcription start sites (TSSs) for the groups in **(b)**. Dotted lines and annotations indicate medians and sample sizes. **(d)** Absolute expression and splicing effect sizes for shared associations at level 2 before and after requiring both QTL types to target the same gene. **(e)** TSS distances for the shared associations in **(d)**. **(f)** Distances from gene-level lead sQTLs to associated donor and acceptor sites for all, sQTL-only, and shared associations at level 1. Donor distances are plotted left of zero and acceptor distances right; annotations give median absolute distances and sample sizes. Throughout, "eQTL–sQTL (Expression)" and "eQTL–sQTL (Splicing)" denote the expression-side and splicing-side associations of variants represented in both traits. Brackets indicate pairwise tests; NS, not significant; $^{**}P < 0.01$; $^{***}P < 0.001$.

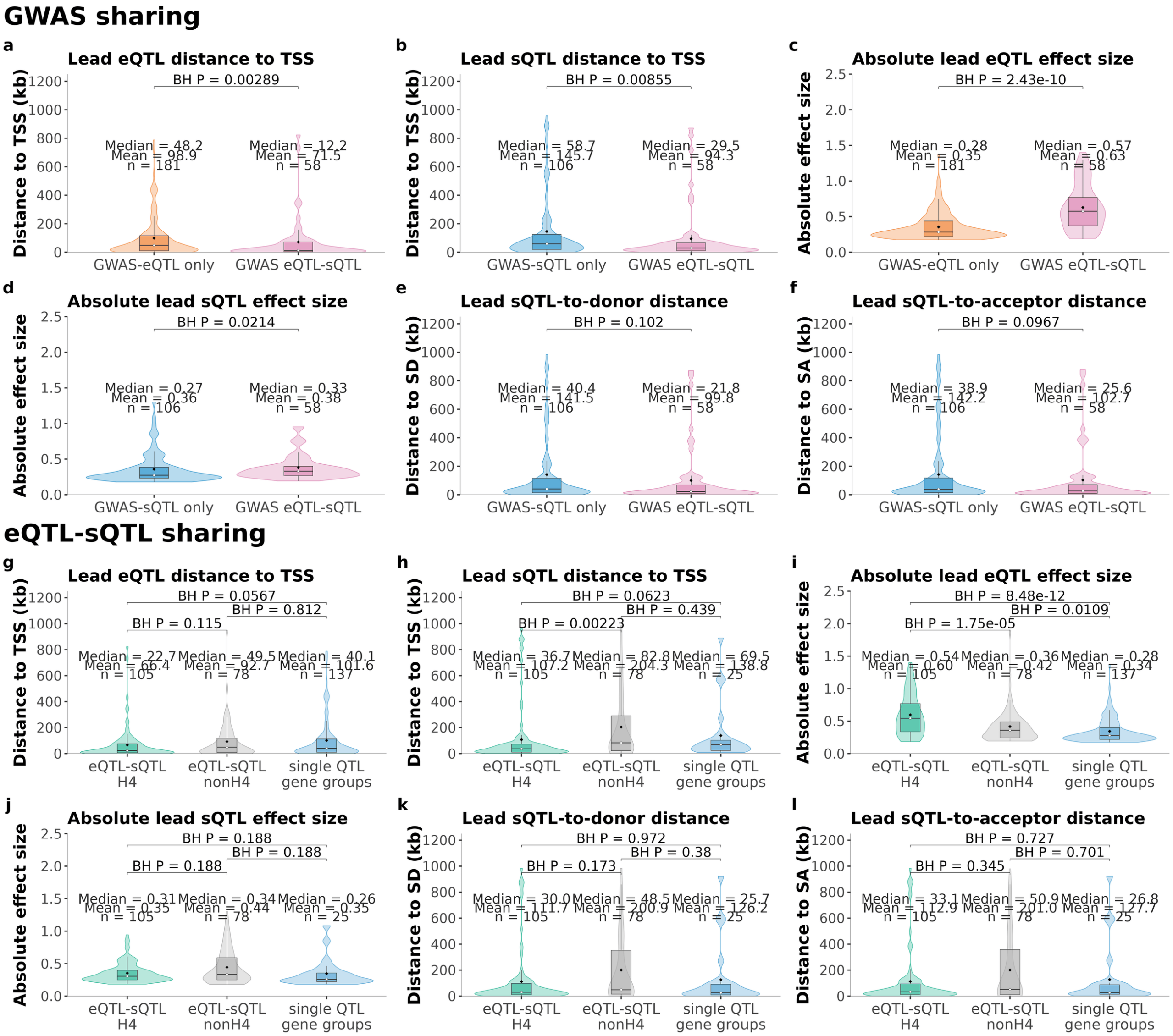


**Supplementary Fig. 24: Lead QTL characteristics stratified by GWAS sharing and eQTL–sQTL sharing.** Distributions of lead eQTL and sQTL distances to the transcription start site (TSS), absolute effect sizes ($|\beta|$), and lead sQTL distances to splice donor (SD) and splice acceptor (SA) sites. Distances are shown in kilobases (kb). **Top: GWAS sharing.** Genes with only an eQTL or only an sQTL colocalized with GWAS are compared with genes for which both molecular traits colocalized with GWAS. **Bottom: eQTL–sQTL sharing.** Genes with evidence of eQTL–sQTL colocalization (H4) are compared with genes with both molecular traits detected but without evidence of colocalization at the specified threshold (nonH4), and genes with only one molecular QTL type detected (single QTL gene groups). Violin plots show the distributions, with embedded box plots indicating medians and interquartile ranges and black diamonds indicating means. Median and mean values and the number of genes included ($n$) are annotated for each group. Brackets indicate pairwise comparisons, with Benjamini–Hochberg-adjusted $P$ values displayed above.
